# Differential effects of immobilized CCL21 and ICAM1 on TILs with distinct expansion properties

**DOI:** 10.1101/2025.03.02.641030

**Authors:** Sofi Yado, Rawan Zoabi, Karin Brezinger-Dayan, Shira Albeck, Tamar Unger, Moran Meiron, Alessio D. Nahmad, Aya Tzur Gilat, Michal J Besser, Benjamin Geiger

**Affiliations:** Department of Immunology and Regenerative Biology, Weizmann Institute of Science, Rehovot, Israel; Samueli Integrative Cancer Pioneering Center, Rabin Medical Center, Petah Tikva, Israel; Department of Life Science Core Facilities, Weizmann Institute of Science, Rehovot, Israel; Orgenesis Biotech, Israel; Dept. of Clinical Microbiology and Immunology and Felsenstein Medical Research Center, School of Medicine, Tel Aviv University, Tel Aviv, Israel; Davidoff Center, Rabin Medical Center, Petah Tikva, Israel

**Keywords:** T cells, Tumor-infiltrating lymphocytes (TILs), TIL expansion, Rapid Expansion Protocol (REP), pre-REP TILs, Adoptive T cell therapy (ACT), CCL21 + ICAM1 synthetic immune niche (SIN), T-cell morphology, Immunophenotyping

## Abstract

Adoptive T cell therapy (ACT), particularly tumor-infiltrating lymphocyte (TIL), holds great promise for cancer treatment, yet it still faces major challenges such as variability in expansion rates, cytotoxic potency and immune suppression. Recent studies suggest that a “synthetic immune niche” (SIN), composed of immobilized CCL21 and ICAM-1, enhances both the expansion and cytotoxicity of murine and patient-derived T cells. Here, we investigate the mechanism underlying the expansion variability by identifying morphological and molecular markers that distinguish low- and high-expanding TILs and predict their *ex vivo* expansion potential. We further developed novel SIN-based strategies that differentially reinforce the efficacy of both low- and high-expanding TILs. We demonstrate that a 14-day REP with feeder cells and SIN facilitates the proliferation of the low-expanding cells, while the high-expanding counterparts benefit from a sequential expansion protocol of 7 days with feeder cells only, followed by 7 days with SIN treatment. At the end of the REP both TIL populations display high levels of granzyme B and perforin and reduced levels of exhaustion markers. In conclusion, our findings demonstrate that the refined CCL21+ICAM1 SIN treatment improves expansion rates and activation profiles of both TIL populations, thereby enabling personalized SIN-enhanced protocols for TIL-based immunotherapy.

## Introduction

Adoptive T cell therapy (ACT) has emerged as a transformative approach to cancer treatment, offering the potential for highly personalized and targeted cure ^1–4^. Among its various strategies, tumor-infiltrating lymphocyte (TIL) therapy stands out due to its broad applicability across a range of solid tumors, including melanoma, breast cancer, cervical cancer, colorectal cancer and non-small cell lung cancer ^5–8^. This approach was first tested in melanoma patients and resulted in an impressive objective response rates of over 50% and a complete remission rate of up to 24% ^9–11^. On February, 2024, the US FDA granted approval for TIL therapy as an autologous cellular treatment for patients with metastatic melanoma ^12^. TIL therapy is based on the isolation and expansion of T cells from a patient’s own tumor specimens and the use of their anti-tumor activity for the elimination of cancer cells in the patient. Tumor fragments are subjected to a pre-Rapid Expansion Protocol (pre-REP), in which a heterogeneous mixture of cells are expanded for 2–3 weeks in culture with interleukin-2 (IL-2)-enriched media, generating predominantly T lymphocytes (pre-REP TILs) which are further expanded for 14 additional days, to yield a clinically-relevant number of cells in a REP process in the presence of IL-2, irradiated peripheral blood mononuclear cells (PBMC, “feeder” cells), and anti-CD3 antibody. Expanded TILs are then harvested and administered back to the lympho-depleted cancer patient ^13,14^.

While there are unique advantages to TIL therapy over single antigen-targeted adoptive cell therapies, such as CAR-T cells, due to the broad antigenic heterogeneity encompassed by TILs and their intrinsic ability to traffic to the tumor site ^15^, there are still major challenges encountered in their wide application. These include variations in the REP expansion yields ^16,17^, immune suppression by checkpoint molecules such as LAG-3, CTLA-4 and PD-1 ^18^; and, often, insufficient numbers of potent autologous T cells that are needed for a clinical scale therapy ^17,19^. Translating these requirements into numbers indicates that for launching a successful TIL-based therapy, the number of potent T cells needed is on the order of 1×10^10^ to 1×10^11^ cells ^20,21^, which requires a major *ex vivo* expansion step on the order of ∼1000-fold or greater.

To address these challenges, we recently developed a 2D stimulatory surfaces that acts as a “synthetic immune niche” (SIN) that supports the *ex vivo* expansion of T cells while maintaining and even enhancing their intrinsic cytotoxic activity. This SIN consisted of tissue culture surfaces, coated with the chemokine C-C motif Ligand 21 (CCL21) and the Intercellular Adhesion Molecule 1 (ICAM-1) ^22,23^. CCL21, secreted by endothelial and stromal cells in the lymph node ^24^, plays key roles in different features of immune responses, including recruitment of T cells and dendritic cells (DCs) ^25,26^, facilitating immune cell migration ^27^, priming of T cells for immune synapse formation ^28^, and co-stimulation of naïve T-cell expansion and Th1 cell polarization ^29–31^. ICAM1 participates in the formation of immune synapses and in the promotion of T-cell activation through binding to its integrin receptor, LFA1 (lymphocyte function-associated antigen 1) ^32^. These two factors were previously shown to act synergistically, as CCL21 increases LFA1 responsiveness to ICAM1, and mediates the arrest of motile lymphocytes on ICAM1-expressing DCs, endothelial cells, and other T cells within their microenvironment ^33,34^.

In our previous studies, we demonstrated that culturing of activated murine CD4^+^ ^23^ or CD8^+ 22^ T cells with CCL21+ICAM1 based SIN significantly increased the expansion of both T-cell subsets. In addition, incubating OVA-specific CD8^+^ T with this SIN increases their efficiency in killing ovalbumin-expressing B16 melanoma cultured target cells, as well as their tumor suppressive activity *in vivo* ^22^. Furthermore, recent study has demonstrated the capacity of a CCL21+ICAM1-based SIN to induce an “optimal interplay” between the expansion and cytotoxicity of CD8^+^ T cells ^35^. More recently, we demonstrated that the CCL21+ICAM1 SIN affects patient-derived TILs and increases their expansion rate as well as the expression of specific activation markers in treated cells ^36^. These findings highlighted the potential of SIN to optimize the clinical utility of TIL therapy. That said, this study also revealed considerable variability in TIL responsiveness to the SIN stimulation during the REP process.

To gain insight into the mechanisms underlying the responsiveness of TILs to the SIN stimulation, we conducted, in the present study, a comprehensive morphological and molecular phenotyping of multiple TILs specimens immediately following the pre-REP stage, and showed that the phenotyping results enable us to predict the level of expansion of each TIL following the REP process. We, then, explored possibilities for enhancing the expansion yields of the different TILs, by stimulating them with the CCL21+ICAM1-based SIN. We show here that the pre-REP early TILs are heterogeneous in their expansion capacity, with two non-overlapping subsets, displaying either low- or high-proliferation properties. These two sub-populations responded differently to the SIN stimulation: the low-expanding TILs underwent major enhancement of their expansion when cultured on the SIN for 2 weeks together with feeder cells, while the high-expanding TILs required a temporal separation between the feeder cell and the SIN treatment. Based on these results, we propose here a novel TIL expansion process, according to which pre-REP TILs will be subjected first to morphometric and multispectral imaging profiling, and those predicted to be a low-expanding TIL, will undergo 14-day long REP in the presence of both feeder cells and SIN, while TILs predicted to be high-responders will be cultured for 7 days with feeder cells only, followed by a 7 days of the CCL21+ICAM1 SIN, in the absence of feeder cells. Our results suggest that this SIN can significantly optimize TIL expansion and reinforce the effectiveness of TIL-based immunotherapy.

## Materials and methods

### Patient-derived tissues and study approval

Tumor tissue samples from six non-small-cell lung cancer (NSCLC) patients (T19, T01074, T16, T20, T93 and T99) and one tumor sample from a bladder cancer patient (T6) were provided by the “Israel National Biobank for Research” (MIDGAM). Additional three tissue samples from renal cell carcinoma (RCC) patients (T160, T55 and T56) were obtained from Samueli Integrative Cancer Pioneering Institute, Rabin Medical Center.

All procedures were conducted in compliance with the declaration of Helsinki (approval no. MID-037-2020). The study was approved and overseen by the Weizmann Institutional Review Board (approval #2358-1).

### Generation and expansion of pre-REP TIL cultures

The establishment of TIL cultures was performed as previously described ^11,37^. Briefly, fragmentation, enzymatic digestion and cell remnants technique were used to isolate TILs from surgically resected lesions. During the first week, non-adherent TILs were transferred to a new 24-well plate, and cultured separately from the adherent cancer cells. TILs were cultured either in RPMI 1640 medium (Biological Industries, Beit Haemek, Israel), supplemented with 10% human AB serum (Gibco, Thermo Fisher Scientific, Waltham, MA, USA), 2mM glutamine (Biological Industries), 1% penicillin–streptomycin (P/S) solution (Biological Industries) and 3,000 IU/ml IL-2 (Akron Biotech, Florida, USA) (” complete medium”), or in X-VIVO medium (Lonza, Basel, Switzerland), supplemented with 50 µg/ml gentamicin (B Braun Medical, Melsungen, Germany) and 3,000 IU/ml IL-2 (Akron Biotech). Cells were split or medium was added every 2 to 3 days, to maintain a cell concentration of 0.5–1.0 × 10^6^ viable cells/ml. TIL cultures were established within 2 to 3 weeks.

### Substrate Functionalization

Substrate functionalization was performed by overnight incubation in phosphate-buffered saline (PBS) with 10 μg/ml CCL21 and 100 μg/ml ICAM1 (produced by the Structural Proteomics Unit, Weizmann Institute, Rehovot, Israel). Detailed information on the protein production and purification can be found in the supplementary information.

### Rapid expansion protocol (REP) with CCL21 and ICAM1 surface coating

Following the establishment of TIL cultures, a standard 14-days REP was initiated by stimulating 10,000 pre-REP TILs with anti-CD3 (OKT-3) antibody (30 ng/ml, MACS GMP CD3 pure; Miltenyi Biotec, Bergisch Gladbach, Germany), 3,000 IU/ml IL-2 (Akron Biotech), and a mixture of irradiated (50 Gy) peripheral blood mononuclear cells (PBMC) feeder cells, obtained from two non-related donors at a 200:1 ratio of feeder cells to T cells. Throughout the REP, TILs were cultured either in complete medium supplemented with 25mM HEPES Buffer (Biological Industries), 50 µg/ml gentamicin (B Braun Medical) and 50 µM β-mercaptoethanol (Biological Industries), pH 7.0, or in X-VIVO medium (Lonza), supplemented with 5% EliteGro™-Adv (Biomedical EliteCell Corp., Beijing, China), 50 µg/ml gentamicin (B Braun Medical) and 3,000 IU/ml IL-2 (Akron Biotech) pH 7.0. REP was performed in 24-well plates and CCL21+ICAM1 coating was prepared one day prior to the initiation or cell splitting. On day 5, 66% of the medium was replaced with fresh medium, irrespective of whether lymphocyte growth was visible. TILs were cultured for 14 days and split on day 7 and 12 to maintain a cell concentration of 0.5-1.0×10^6^/ml. Alternatively, pre-REP TILs were expanded following the same REP procedure, using a medium containing 66% of conditioned medium (CM) of feeder cells cultured for 7 days without TILs (2×10^6^ cells/well). The number of viable TILs was determined at day 14 using a CellDrop™ FL automated cell counter device (DeNovix, Wilmington, DE, USA) and trypan blue staining solution. At the end of the REP, cells were analyzed phenotypically by FACS. A schematic representation of REP process with CCL21 and ICAM1 surface coating is shown in **Supplementary Figure 1**.

### Spectral flow cytometry

Pre-REP TILs underwent a 14-day REP with or without substrate coated SIN as described above. Surface and intracellular marker expression were then tested by spectral flow cytometry. Brefeldin A (5 μg/ml, BioLegend, San Diego, CA, USA) was added to the cultures during the last 4 h of incubation to allow intracellular cytokine accumulation. Expanded TILs were then detached from the substrate, placed in a U-shaped 96-well plate, and washed with PBS + 3% FBS (Biological Industries, “washing buffer”). The supernatant was aspirated, and the cells were surface stained (at room temperature for 30 min) with LIVE/DEAD Fixable Blue dead cell stain (1:1000; Invitrogen, Paisley, UK) and with the following fluorescent monoclonal antibodies from the following sources: CD3-Alexa 700, CD56-FITC, CD4-PerCP, CD8-BV510, CD25-APC-Fire 810, CD69-BV750, PD-1-BV421, LAG-3-BV650, CD197-BV711, CD45RO-BV605 and TCR γ/δ-PE [all from BioLegend]. The cells were then washed, fixed and permeabilized [BioLegend] and stained (at room temperature for 30 min) intracellularly with antibodies from the following sources: Granzyme B-Alexa 647, Perforin-Alexa 594 and IFNγ-APC [all from BioLegend]. After washing, the data were acquired using an Aurora (Cytek, CA, USA) spectral flow cytometer, and the data were analyzed using FlowJo software (Ashland, OR, USA).

### Microscopy and image analysis

Pre-REP TILs were stimulated and cultured for 14 days with or without substrate coated SIN as described above. The morphology and growth patterns of the TILs were examined using a Celldiscoverer 7 microscope (Carl Zeiss Ltd.) equipped with a Plan-Apochromat 20x/0.7 and a 1x Tubelens connected to an Axiocam 702 camera (Carl Zeiss Ltd.). Images were taken on day 0 (end of pre-REP stage) and 14, using oblique illumination, and analyzed using ImageJ software (https://imagej.nih.gov/ij/). Pre-REP TILs were segmented using Cellpose algorithm ^38^, and their projected area and circularity were measured using ImageJ software.

### Proteome Profiler Array

The Proteome Profiler Human Cytokine Array Kit (R&D Systems, Minneapolis, MN, USA) were utilized to identify cytokines and chemokines secreted by feeder or TIL cells in response to SIN stimulation. A total of 36 cytokines were analyzed. In short, supernatants were collected at day 5 of the REP from feeder cells cultured alone or low- and high-expanding TIL cultures stimulated with feeder cells. Array membranes (consisting of capture antibodies spotted in duplicate on nitrocellulose) were incubated with 0.5 mL supernatant and soluble proteome was analyzed following the manufacturer’s instructions using the Bio-Rad ChemiDoc Touch Imaging System (Bio-Rad, Hercules, CA, USA). The spot pixel density was measured by ImageJ analysis software in arbitrary units.

### Statistical analysis

The significance of variation between groups was evaluated using a non-parametric two-tailed Student’s t-test.

## Results

### Heterogeneity of the proliferative potency of pre-REP TILs

In the clinical setting, the standard process for TIL expansion consists of two major steps; a pre-REP phase that includes the extraction and expansion of TILs from tumor fragments in the presence of high concentration of IL-2, followed by a REP process that involves expansion of the extracted TILs in culture supplemented with IL-2, anti-CD3 mAb (OKT-3), and irradiated PBMC that serve as feeder cells. Following the REP process, the cells are harvested and prepared for infusion back into the patient ^39^. Naturally, given the intrinsic patient-to-patient variation, and the effects of specific anti-cancer treatment history, the differences in the expansion rates and in the efficacy of the TIL therapy are highly variable ^40^.

To explore the heterogeneity in the intrinsic expansion profile of pre-REP TILs, we conducted a systematic REP screen of 10 pre-REP samples derived from lung, kidney, bladder and endometrial tumors (see Materials and Methods section), that were expanded with diverse feeder cells (F_1,_ F_2_ and F_3_, each prepared by mixing cells derived from two non-related healthy donors). The expansion was calculated based on the fold change in cell number at the end of the REP (day 14) compared to the number of TIL at initiation (at day 0). Altogether, a total of 17 different combinations of TILs and feeder cells were tested (**Figure 1a**’ and **a’’**).

**Fig. 1:**
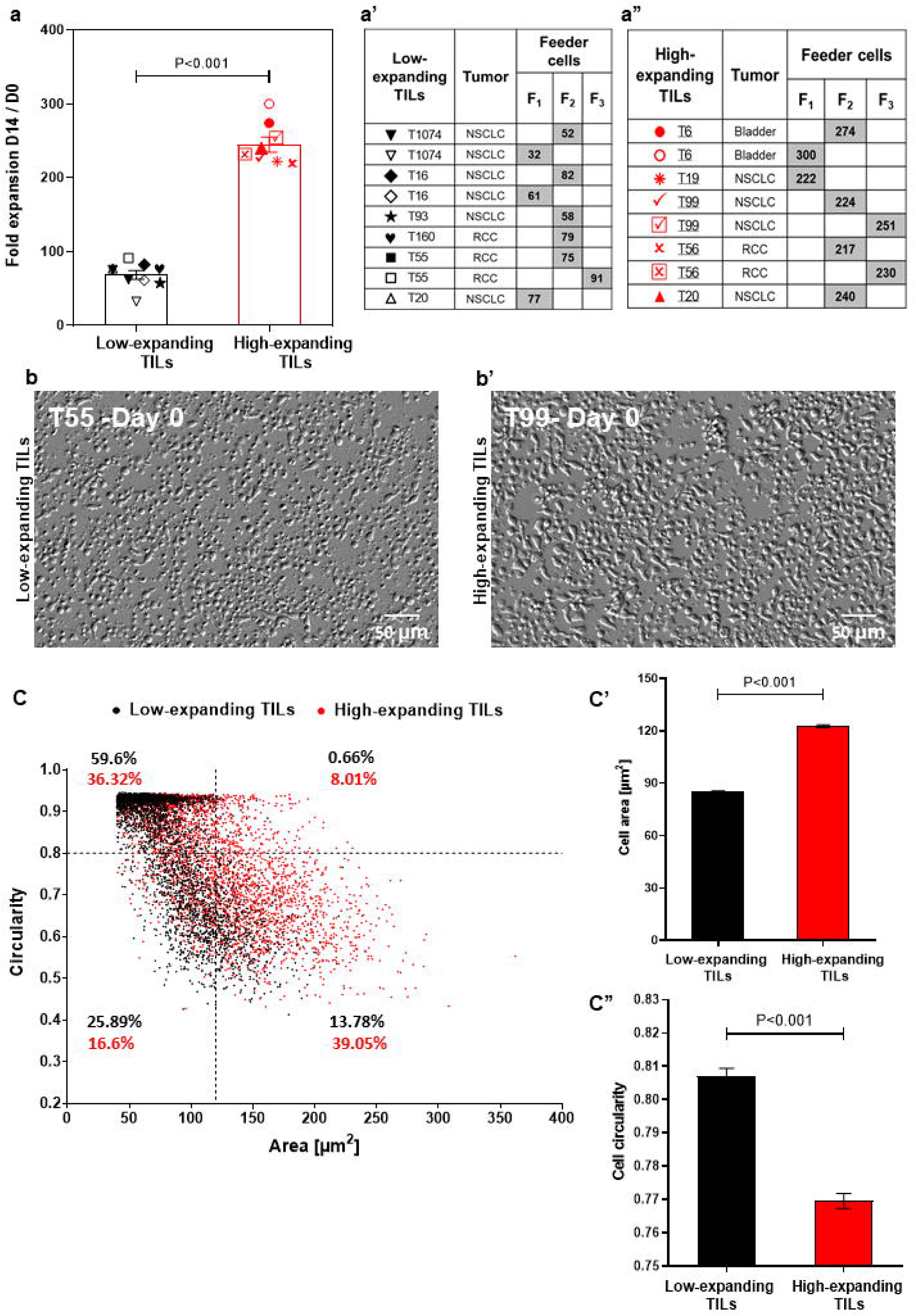
Heterogeneity of the intrinsic expansion and morphological profiles of pre-REP TILs. **(a)** Fold expansion at day 14 of the REP for different TIL-feeder combinations (n_TIL_=10; n_feeder_=3). The feeder-TIL pairs are divided into two groups: “high-expanding TILs” (expansion >100 fold, red symbols) and “low-expanding TILs” (expansion <100 fold, black symbols). Feeder cells (F_1_, F_2_, and F_3_) were prepared by mixing cells derived from two non-related healthy donors. **(a’)** Pairing scheme of the low-expanding TILs and different feeder cells (indicated by the gray boxes). **(a”)** Pairing scheme of high-expanding TILs and different feeder cells (indicated by the gray boxes). Data in **(a)** are shown as mean ± SEM of three independent experiments. Calculated p-values (using standard t-test) are as indicated in the figure. (**b** and **b’)** Representative images of low-expanding **(b)** and high-expanding **(b’)** pre-REP TILs that depict morphological differences between the cells based on oblique illumination-based microscopy. Scale bar: 50 μm. **(c)** Morphological profiling of the low- and high-expanding TILs shown in b and b’ (T55 and T99 respectively), based on their projected area and circularity. The relative prominence of high- and low-expanding TILs in this scatter-plot indicates that the low-expanding cells are the most prominent population in the small (< 120 µm^2^) and circular (circularity >0.8) cells. The high-expanding cells were the most prominent population among the large (> 120 µm^2^) and polarized (circularity <0.8) cells. **(c’** and **c’’**) The average projected area (**c’)** and circularity **(c’’)** of the low-expanding (black bars) and high-expanding (red bars) cells.

Notably, the TIL expansion process conducted in this study was performed in open multi-well plates, rather than in closed bioreactor or other scaled up systems, and the typical fold expansion was in the range of up to 200-300. That said, comparison of the expansion capacities of 10 pre-REP TIL samples, revealed two, non-overlapping subsets of pre-REP TILs: low-expanding TILs in which the fold change was lower than 100 (average 68.3±5.8) and high-expanding TILs in which the fold change was higher (average 244.8±10.2; **Figure 1a)**. Interestingly, when the same TILs underwent REP in the presence of different feeder cell pairs, different fold expansion values were obtained. Thus, TILs from donor T20 stimulated with feeder cell mix F_1_ showed lower expansion rates compared to those stimulated with feeder cell mix F_2_ (**Figure 1a**), suggesting that the variations between different feeder cells may affect the expansion capacity of TILs. Additionally, light microscopy-based morphometry indicated that at the end of the pre-REP, the high-expanding TILs display higher projected area (122.7±0.85 μm^2^) and lower circularity (0.76±0.002), than the low-expanding TILs (85.14±0.56 μm^2^ and 0.8±0.002, respectively, **Figure 1b** and **b’**). Overall, these results indicate that pre-REP TILs may vary in their intrinsic expansion potential, as well as in their capacity to respond to different feeder cell, which certainly may affect their applicability in therapy.

### Molecular and morphological phenotyping of pre-REP TILs enables a differential prediction of their expansion potency

To gain an insight into the differential properties of the low- and high-expanding TILs, we conducted a retrospective morphological and molecular phenotyping of each of the pre-REP TIL samples. In accordance with our microscopic observations (**Figure 1b** and **b’)**, flow cytometry analysis showed that the high-expanding TILs display significantly higher FSC **(Figure 2a** and **b)** and SSC **(Figure 2a’**and **b’)** intensities, compared to low-expanding TILs, indicating that decreased cell sizes and lower cytoplasmic granularity correlate with low expansion potency.

**Fig. 2:**
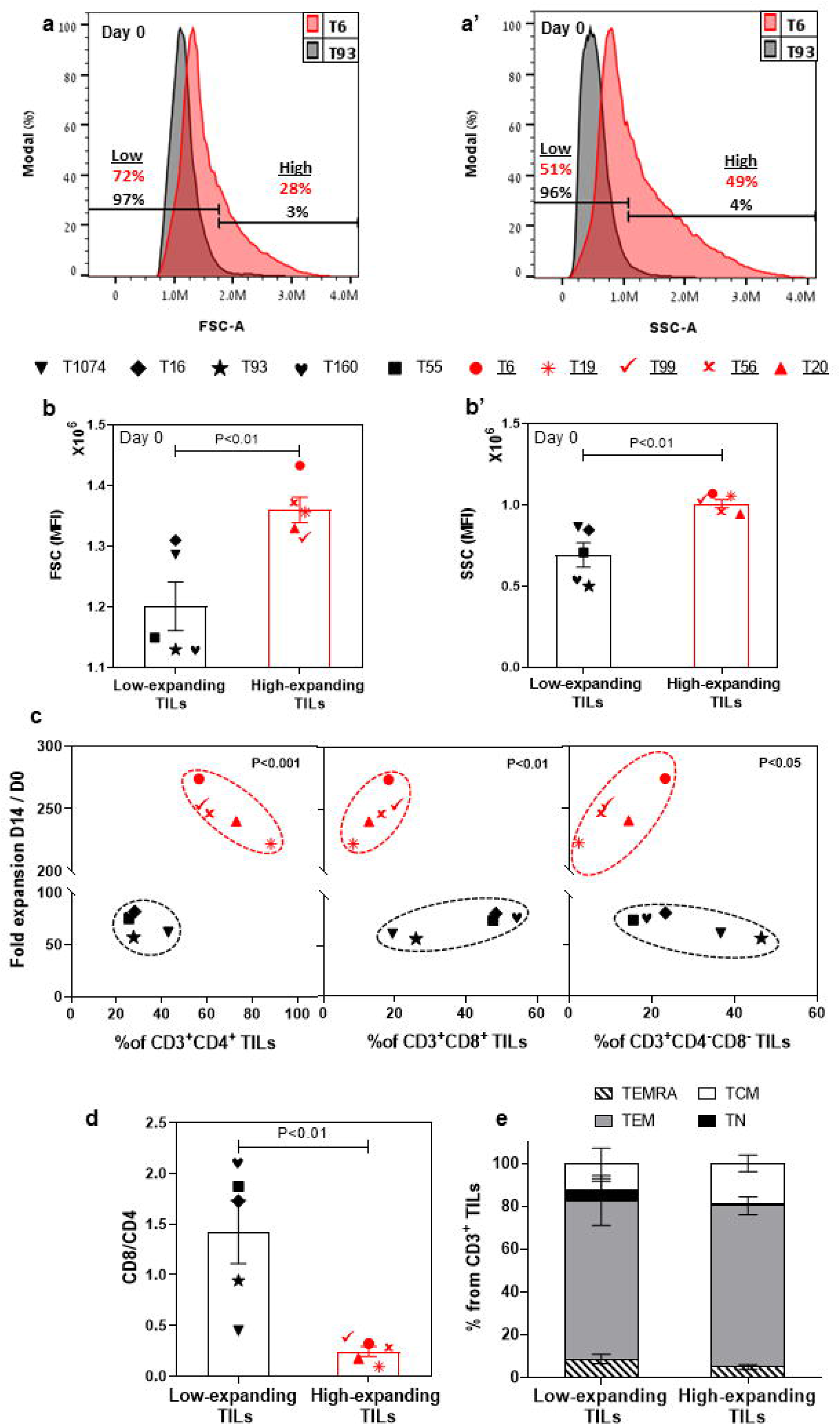
Molecular and morphometric phenotyping of pre-REP TILs enables a differential prediction of their expansion potency. Flow cytometry-based phenotyping of pre-REP samples (n=10). Gating strategy is shown in **Supplementary Fig. 2**. Representative FSC **(a)** and SSC **(a’)** profiles of high- and low-expanding pre-REP TILs (T6 and T93, respectively). (**b** and **b’)** Bar graphs showing the median fluorescence intensity (MFI) of FSC and SSC, respectively, obtained for 10 independent TILs. Data are shown as mean ± SEM of three independent experiments. Calculated p-values (using standard t-test) are as indicated in the figure. (**c)** Fold expansion as a function of mean frequencies of CD4^+^, CD8^+^ and CD4^−^ CD8^−^ T cells within the CD3^+^ lymphocyte population in pre-REP TILs. Calculated p-values (using standard t-test) are as indicated in the figure. Data shown here are representatives of three independent experiments. (**d)** CD8/CD4 ratio in pre-REP TILs. Data are shown as mean ± SEM of three independent experiments. Calculated and p-values (using standard t-test) are as indicated in the figure. (**e)** Distribution of subsets of differentiation markers in pre-REP TILs. The percentages (mean ± SEM) of naïve (TN, CD45RO^+^CCR7^−^), central memory (TCM, CD45RO^−^CCR7^+^), effector memory (TEM, CD45RO^−^CCR7^−^) and effector (TEMRA, CD45RO^+^CCR7^−^) T cells in CD3 population are shown. Data presented here are representatives of three independent experiments.

Next, we evaluated, by flow cytometry, the prominence of specific T cell subsets within the low- and high-expanding CD3^+^ T-cell populations, as well as the respective levels of expression of phenotypic markers on the cells. Surprisingly, this analysis showed that a low frequency of CD8^+^ T cells and high prominence of CD4^+^ T cells (i.e. a lower CD8/CD4 ratio) in the pre-REP TILs correlates with the high proliferation levels of these cells **(Figure 2c** and **d)**, suggesting that this variables may be considered as a conclusive predictive biomarkers for non-responsiveness. It is noteworthy that it had been shown that at the conclusion of the REP expansion, the cells display high CD8^+^ and low CD4^+^ levels (**Figure 7e**, see below). Additionally, we found that the prominence of CD4^−^ CD8^−^ population was greater for low-expanding TILs than for high-expanding TILs **(Figure 2c**, see discussion).

We further characterized the phenotype of the pre-REP TIL samples using selected representative markers that are known to be involved in T-cell activation (CD25, CD69), exhaustion (PD-1, LAG-3), cytotoxic function (granzyme B, perforin) and overall differentiation (CD45RO, CCR7) of T cells. This analysis indicated that within the high-expanding TILs, the expression of CD25 (a late activation marker) was significantly higher on both CD8^+^ **(Supplementary Figure 3a)** and CD4^+^ **(Supplementary Figure 3c)** T cells, compared to their levels on the low-expanding TILs, whereas no significant difference was evident in the level of expression of CD69 (an early activation marker, **Supplementary Figure 3b and d)**. Together, these results indicate an efficient and sustained activation of the highly proliferative TILs.

Previous studies reported that immune checkpoint inhibitory receptors can alter T-cell function and expansion both *in vitro* and in *in vivo* ^41^. To address the possible involvement of the checkpoint inhibitory system in the REP expansion, we checked the expression levels of two inhibitory receptors: PD-1 and LAG-3 in pre-REP TILs. We noted that among the high-expanding TILs, the expression of LAG-3 was significantly lower on both CD8^+^ **(Supplementary Figure 3e**, *p*<0.05) and CD4^+^ **(Supplementary Figure 3g**, *p*<0.05) T cells, compared to the low-expanding TILs, while no significant change was evident in the expression level of PD-1 (**Supplementary Figure 3f and h**). Given that LAG-3 is associated with reduced proliferative capacity ^42,43^, his relatively high expression within low-expanding TILs could account for the low expansion potential of these TILs.

Additionally, we determined the functional state of pre-REP T cells by analyzing granzyme B **(Supplementary Figure 3i)** and perforin **(Supplementary Figure 3j)** differential expression on the high- vs. low-expending CD8^+^ T cells. Surprisingly, we found that the expression of granzyme B and perforin were essentially the same on CD8^+^ T cells in the low-and high-expanding cells. Finally, we tested the differentiation status of pre-REP TILs based on the expression of CD45RO, in combination with CCR7. As expected, we found that both low- and high-expanding TILs were mostly effector memory T cells (TEM: CD45RO^+^CCR7^−^, **Figure 2e**). Collectively, the cellular morphology and molecular phenotyping of pre-REP TILs, reveals significant differences that enable clear prediction of their REP expansion capacities.

### The differential effects of SIN treatment on low- and high-expanding TILs during the REP process

The use of a SIN, consisting of immobilized CCL21 and ICAM1 for enhancing TIL-based cancer therapy was motivated by the limited efficacy of the current TIL-based therapy ^44–46^, and the apparent capacity of the CCL21-ICAM1 SIN to stimulate the proliferation of cytotoxic T cells, while maintaining, or even enhancing their cytotoxic potency ^22,35^. To determine the differential effects of SIN treatment on the high- and low-expanding TILs, we have exposed both populations of cells to REP process for 14 days in 24-well plates, either coated with CCL21 + ICAM1 or uncoated.

As presented in **Figure 3a**, five out of nine low-expanding TIL cultures tested showed higher expansion values following exposure to CCL21+ICAM1 SIN throughout the 14-day REP. In these cultures (for details, see **Figure 3e** and **e’**), the expansion of SIN-treated TILs was, on average, 4.8 ± 0.4-fold greater (**Figure 3b)**, reaching an overall 344.3 ± 51.5-fold expansion (relative to the number of cells on day 0) for SIN-treated TILs, compared with 71.0 ± 7.1-fold expansion for the same TILs, cultured on uncoated surfaces.

**Fig. 3:**
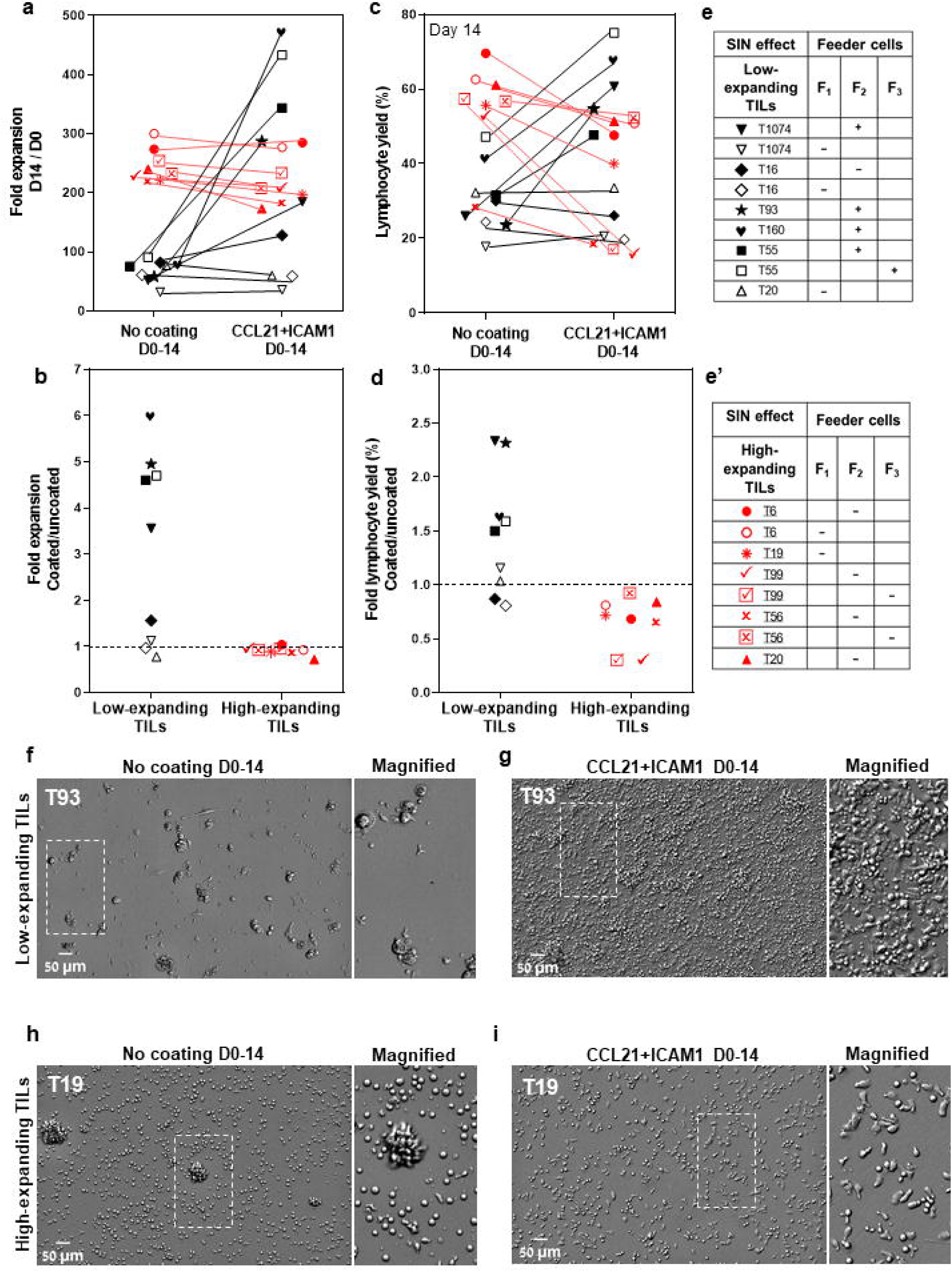
The differential effects of CCL21+ICAM1 SIN treatment on low- and high-expanding TILs during the REP process. (**a** and **c)** Fold expansion (relative to the relevant numbers on day 0) **(a)** and lymphocyte yield (assessed by flow cytometry, at day 14 of the REP) (**c**) of TILs at day 14 for different TIL-feeder combinations (n_TIL_=10; n_feeder_=3), in the presence or absence of CCL21+ICAM1 SIN stimulation. (**b)** and **(d)** show the fold expansion and lymphocyte yield ratios of T-cells cultured on SIN-coated surfaces compared to those cultured on uncoated surfaces. Data are shown as the mean of three independent experiments. Tables **e** and **e’** mark the SIN effect on the expansion of the low- and high-expanding TILs, respectively. (+) sign indicates a SIN-induced increase in TIL expansion, and the (−) sign represents a lack of effect or even suppression of expansion. **f-i,** Representative images of low- **(f** and **g)** and high-expanding TILs **(h** and **i)**, grown for 14 days either on the uncoated substrate **(f** and **h)** or on the substrate-immobilized CCL21 + ICAM1 **(g** and **i)**, demonstrating a prominent SIN-induced increase in the expansion of most of the low-expanding TILs **(g)** compared to those growing on uncoated substrates **(f)**. On the other hand, SIN treatment of the high-expanding TILs revealed an unexpected loss of stimulatory effect of the SIN, or even suppression of TIL expansion, pointing to an apparent strong conflict between the feeder cell stimulation and and the concimitent SIN stimulation, manifested by a smaller yield of these cells **(i)**, compared to untreated cells **(h)**. Magnified views of the marked areas are shown on the right of each image. Scale bars: 50 μm.

Moreover, the lymphocyte yield (displaying >90% viability) among these TIL cultures was higher for cells cultured on CCL21 + ICAM1 substrates than for cells cultured on uncoated substrates **(Figure 3c** and **d)**. Notably, this level of expansion was comparable to, or even higher than that of the high-expanding TILs, cultured on uncoated surfaces **(Figure 3a)**. Unexpectedly, all the high-expanding TIL cultures, as well as one low-expanding TIL (T16) that displayed exceptionally low fold expansion values, failed to respond to the REP process following the CCL21+ICAM1 SIN treatment **(Figure 3a-d)**, indicated by a decrease of fold expansion values by 11.0±3.2% (**Figure 3a, b),** and a decrease of the lymphocyte yield among these TIL cultures by 33.7±8.6% (**Figure 3c, d)** in the presence of 2 distinct feeder cells.

Further insight into the differential cellular effects of SIN treatment on low- and high-expanding TILs at the REP endpoint was obtained by microscopy-based imaging, which indicated that stimulation of low-expanding TILs with the SIN yielded significantly higher numbers of large and polarized T cells (**Figure 3g)**, compared to untreated cells (**Figure 3f)**. On the other hand, stimulation of the high-expanding TILs with the SIN decreased the yield of T cells, however the prominence of large and polarized T cells was increased (**Figure 3i)**, compared to untreated cells (**Figure 3h)**.

Taken together, these results show that SIN treatment has radically different effects on low-expanding cells (whose proliferation is augmented) and high-expanding cells, whose proliferation is, apparently suppressed.

### The tri-partite interplay between TILs, feeder cells and the CCL21+ICAM1 SIN during REP

The results described above revealed an apparently complex interplay between an essential component of the REP process, namely the feeder cells and the SIN treatment. Specifically, we found that feeder cells are essential for successful REP and cannot be replaced by the CCL21+ICAM1 SIN alone (**Figure 4a** and **a’)**, yet simultaneous exposure of low- and high-expanding TILs to both feeder cells and SIN throughout the REP process, leads to radically different outcome; low-expanding TILs display an enhanced expansion, while the high-expanding TILs are strongly suppressed by the same treatment. To explore this intriguing interplay between the feeder cells and the SIN, we tested three aspects of the feeder-cells’ effect on the REP process: (i), Does the REP efficacy depend on the presence of live feeder cells in the TIL culture? (ii), Similarly, is the apparent feeder-SIN conflict in the case of the high-expanding TILs dependent on feeder cells-TIL interaction? (iii), Which component of the SIN (CCL21 or ICAM1) is responsible for the apparent conflict with the feeder cells?

**Fig. 4:**
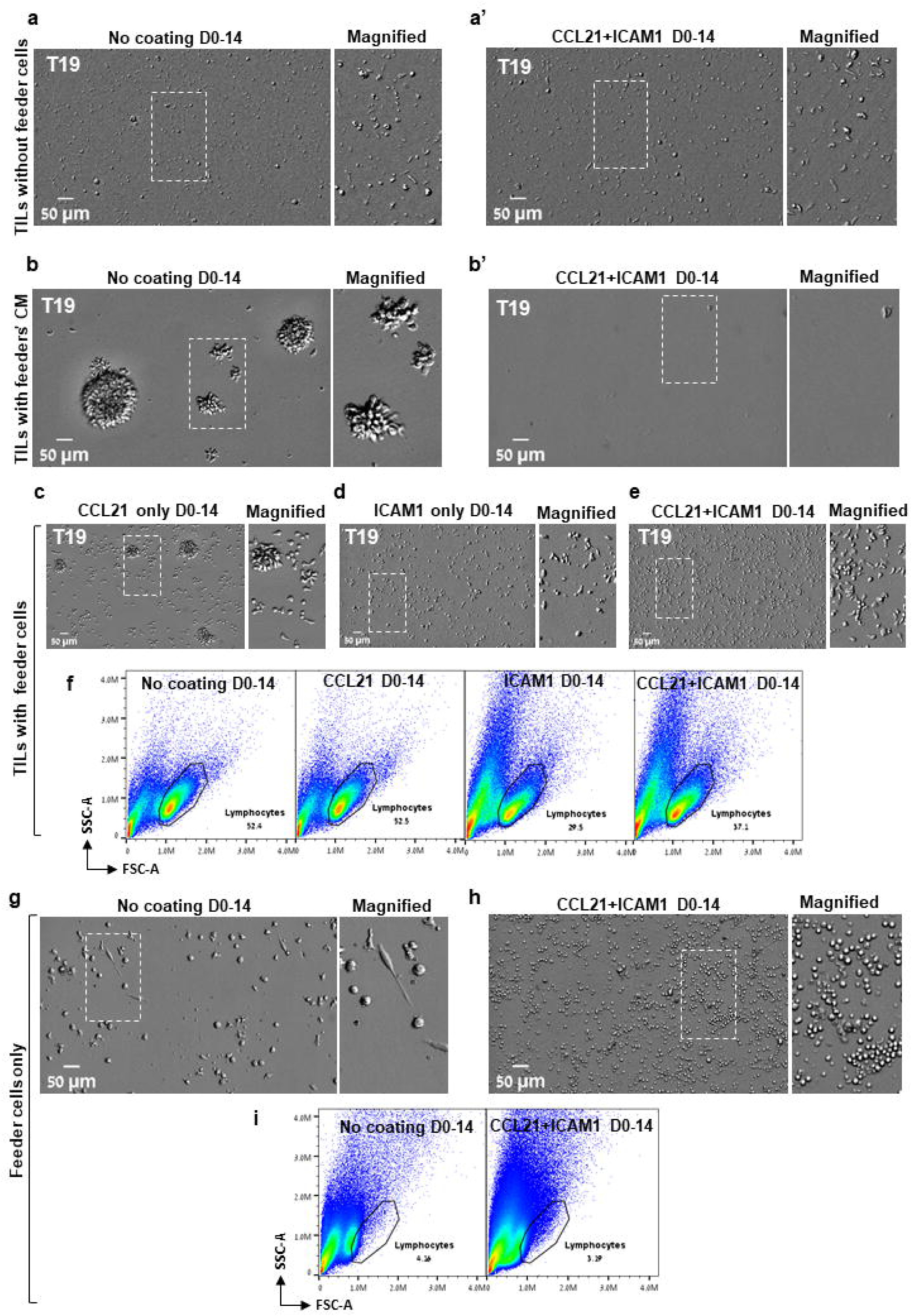
The tri-partite interplay between TILs, feeder cells, and the CCL21+ICAM1 SIN during the REP. **(a-b’)** Representative images of high-expanding TILs cultured for 14 days without feeder cells **(a** and **a’**) or with feeder cells’ conditioned medium (CM, **b** and **b’**) on either an uncoated substrate **(a** and **b)** or on substrate-immobilized CCL21 and ICAM1 **(a’** and **b’)**. **(c-e)** Representative images of high-expanding TILs cultured for 14 days with feeder cells on a substrate coated with the individual SIN components, namely CCL21 **(c)**, ICAM1 **(d)**, or both CCL21 and ICAM1 **(e)**. Magnified views of the marked areas are shown on the right of each image. Scale bars: 50 μm. (**f)** Representative flow cytometry analyses of high-expanding TILs stimulated for 14 days on an uncoated substrate or on substrates coated with the SIN components, namely CCL21, ICAM1, or both, showing the TILs’ forward and side scatter profiles. (**g** and **h)** Representative images of feeder cells (without TILs) cultured for 14 days on either an uncoated substrate **(g)** or on substrate-immobilized CCL21 + ICAM1 **(h).** Magnified views of the marked areas are shown on the right of each image. (**i)** Representative flow cytometry analysis of feeder cells, cultured for 14 days on either an uncoated substrate or on substrate-immobilized CCL21 + ICAM1. The lymphocytes were determined by the forward and side scatter profiles. Experiments were conducted with three TILs-feeders combinations, namely T19/F_1_, T99/F_2_ and T6/F_2_.

To address these questions, high-expanding pre-REP TILs were cultured for 14 days following the preREP in the presence of feeder cells conditioned medium (fcCM), on either uncoated substrate or on substrate-immobilized CCL21 + ICAM1 (for details, see Materials and Methods section). The results shown in **Figure 4b** indicated that the cell-free fcCM supported TIL expansion in the absence of SIN; however, the yield was approximately 4-fold lower compared to irradiated feeder cells (Figure 3h). When SIN was present, fcCM suppressed TIL expansion **(Figure 4b** and **b’)**, similar to its effect in the presence of irradiated feeder cells **(Figure 3i)**. These results suggest that both the support of expansion (in the absence of SIN) and its suppression (in the presence of SIN) are mediated via components released by the irradiated feeder cells to the culture medium. Finally, to determine which of the SIN components is responsible for the feeder cells/SIN conflict, pre-REP TILs were expanded according to the REP process with irradiated feeder cells on either CCL21 alone or ICAM1 alone. Our data revealed that compared to treatment with both CCL21+ICAM1 **(Figure 4e)**, stimulation with immobilized CCL21 alone induced expansion of TILs **(Figure 4c)**, while immobilized ICAM1 treatment suppressed it **(Figure 4d)**. Moreover, as shown in **Figure 4f**, the lymphocyte yield among these TIL cultures was lower for cells cultured on ICAM1 substrates than for cells cultured on CCL21. Interestingly, culturing feeder cells alone for 14 days on SIN-coated and uncoated surfaces indicated that treatment with the SIN decrease the viability of feeder the cells **(Figures 4g-i)**. Taken together, we found that the feeder cells/SIN conflict is affected by three main factors: TILs responsiveness to the REP procedure (“High” vs. “Low” responsiveness), feeder cells’ variability and the presence of surface-immobilized ICAM1.

Given that the feeder cells’ conditioned medium stimulates the proliferation of high-expanding TILs in the absence of SIN **(Figure 4b)**, but this effect is suppressed when SIN is present **(Figure 4b’)**, we evaluated the effects of SIN treatment on the production of diverse cytokines by feeder cells alone as well as by TIL-Feeder co-cultures. To profile these cytokines, we used a cytokine array, measuring the levels of 36 different cytokines and chemokines in the culture medium of representative feeder cells (F_2_ and F_3_) and in feeder-TIL co-cultures (T99/F_2_, T99/F_3_, T55/F_2_ and T55/F_3_) in the presence and absence of the SIN, respectively. As shown in **Figure 5a** and **b**, SIN treatment led to substantial changes in the cytokine profile, particularly within the C-C and C-X-C chemokine ligand families, as well as interleukins. Specifically, SIN treatment induced a major increase in the level of CCL2 in feeder-only cultures (F_2_), along with more modest increase in the levels of CCL3/4 and CXCL1, low increase in IL-6, and low decrease in CXCL10. Analysis of the cytokine profile of the Feeder-TIL co-culture medium from low and high expanding TILs (T99/F_2_ and T55/F_2_) revealed a largely similar increase in the levels of CCL3/4, CXCL1, IL-1β, and IL-6 in response to SIN treatment. In contrast, SIN induced a major reduction in the levels of IL-8 and CXCL10, which are known to drive T cell dysfunction or exhaustion through excessive signaling in various pathological contexts ^47,48^. Given that IL-6 and IL-1β synergistically promote T-cell proliferation and survival ^49,50^, these finding suggest that SIN treatment may enhance T-cell activation and persistence. It is notable that while the co-cultures containing low- and high-expanding TILs displayed a largely comparable response to the SIN treatment, the extent of the effect was often variable (e.g., CCL3/4, IL-2). Comparison of the SIN effect on the cytokine profile in the feeder-TIL co-culture and the feeder-only showed some similarity (e.g., CCL3/4, CXCL1, IL-6), but also major differences, such as the prominence of CCL2 in the feeder-only cultures compared to its modest suppression in the co-culture setting, and the robust SIN-induced suppression of IL-8 in the feeder-TIL cultures, compared to its modest increase in the feeder-only culture. Notably, ICAM1 levels were increased in SIN-treated cells compared with untreated cells across both feeder-only cultures and TIL-Feeder co-cultures, possibly due to leakage from the culture plate. Since CCL3/4 and CXCL1 synergize with ICAM1 to promote the recruitment and activation of T cells during immune responses ^51,52^, their increased secretion following SIN treatment (**Figure 5a** and **b**) could potentially modulate the tumor immune microenvironment, thereby enhancing anti-tumor immunity. A similar cytokine shift was observed in feeder F_3_ cultures (**Supplementary Figure 4a and b**). Collectively, these findings suggest that SIN treatment alters the cytokine milieu in a way that could modulate the tumor immune microenvironment, potentially boosting immune cell activity against the tumor.

**Fig. 5:**
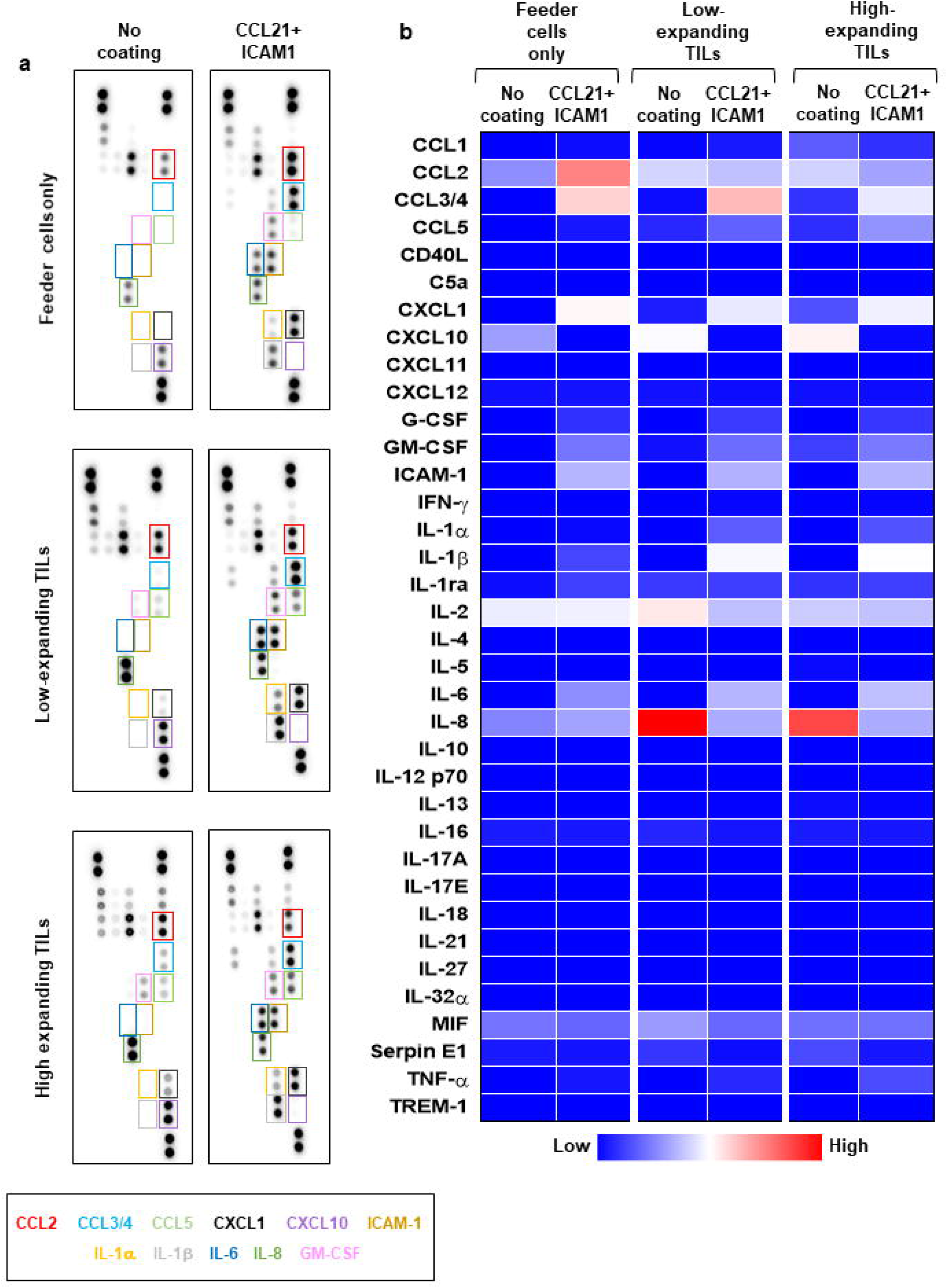
The differential effects of CCL21+ICAM1 SIN treatment on cytokine secretion by low- and high-expanding TILs. Cytokine array analysis of culture supernatants collected at day 5 of the REP from feeder cells cultured alone (F_2_) or low- and high-expanding TIL cultures stimulated with feeder cells (T99/F_2_ and T55/F_2_). **(a)** Representative images for cytokine array panels. **(b)** Quantification of results from the cytokine array analysis. Heatmap represents average values from two dots in the array. The differentially secreted cytokines are highlighted in boxes, and their labels are shown below with corresponding colors. Results of a repeated analysis with feeder T3 are shown in Supplementary Figure 4.

### Temporally separating the feeder and SIN stimulation eliminates the suppressive effect of SIN on the high-expanding TILs, and greatly enhances the overall expansion

The apparent conflict between the simultaneous stimulation of the high-expanding TILs by both the feeder cells and the CCL21+CAM1 SIN raised the question of whether a temporal separation of the two treatments could rescue the potential synergy between them. To test this possibility, we divided the REP into two periods, one week each, with non-overlapping feeder cells and SIN treatments. In a preliminary set of experiments, we tested the optimal order of the SIN versus the feeder cell treatments. These experiments indicated that starting with the SIN stimulation does not support cell survival while co-culturing the TILs with the feeder cells for one week, followed by one week with the SIN only, has a remarkable beneficial effect on TIL expansion.

As shown in **Figure 6a**, this 2-stage stimulation resulted in 2.3 ± 0.1-fold expansion higher than that of cells cultured on uncoated surfaces (**Figure 6a)**. Specifically, the cells reached an overall 524.5 ± 44.8-fold expansion (relative to the number of cells on day 0) for SIN-treated TILs, compared with 244.7 ± 10.2-fold expansion for the same TILs, cultured on days 0-14 on uncoated surfaces (*p*<0.001). Moreover, the lymphocyte yield among these TIL cultures was higher for cells cultured on CCL21 + ICAM1 substrates on days 7-14 than for cells cultured on uncoated substrates on day 0-14 (*p*<0.01) **(Figure 6b)**. In addition, high-expanding TILs stimulated with the SIN on day 7-14 appeared more elongated, with high projected area **(Figure 6d)**, compared to untreated high-expanding TILs, that remain small and round **(Figure 6c)**. Notably, low-expanding TILs do not respond well to a protocol based on one-week co-culture with irradiated feeder cell, followed by one week with the SIN only **(Figure 6e** and **f)**.

**Fig. 6:**
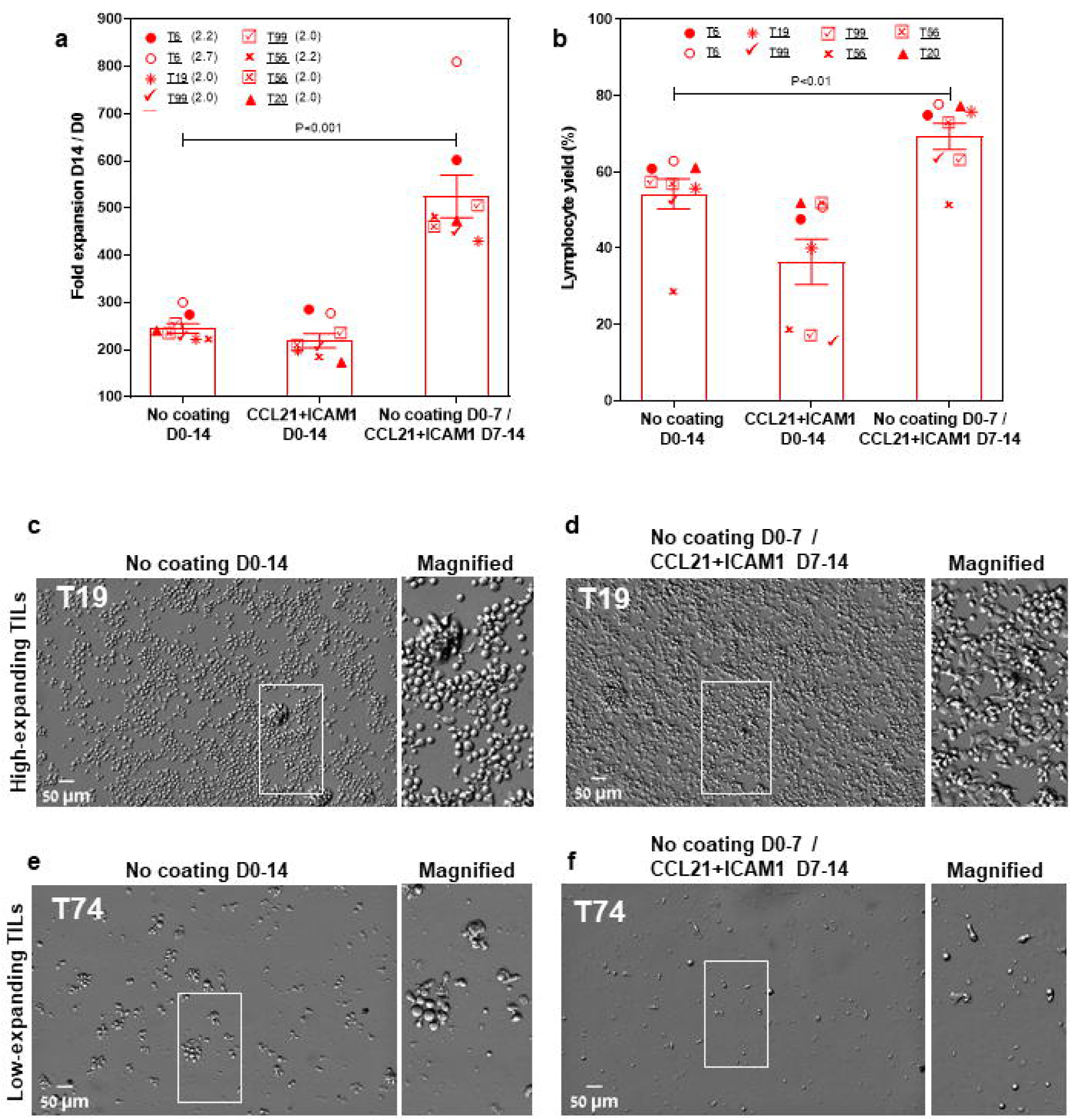
Temporally separating the feeder and SIN stimulations eliminates the suppressive effect of SIN on the high-expanding TILs and greatly enhances the overall expansion levels. (**a** and **b)** Fold expansion (relative to the number of cells on day 0): **(a)** and lymphocytes yield **(b)** (assessed by flow cytometry, at day 14 of the REP) for high-expanding TIL cultures (n=9). The TILs were stimulated for one week with feeder cells only, then washed and incubated for one week with the CCL21-ICAM SIN without feeder cells. (**c-f)** Representative images of high- **(c** and **d)** and low-expanding TILs **(e** and **f)** stimulated for one week with feeder cells, followed by one week with the SIN only. Magnified views of the marked areas are shown on the right of each image. Scale bars: 50 μm.

### Molecular phenotyping of SIN-stimulated TILs at the end of the REP process (day 14)

As shown above, the low-expanding TILs can be treated simultaneously with feeder cells and SIN for 14 days, effectively converting them to high-expanding TILs. The high-expanding TILs, on the other hand, are sensitive to the concomitant stimulation with feeders and SIN, but they respond well to a protocol based on one-week co-culture with irradiated feeder cells, followed by one week with the SIN only.

To explore the impact of CCL21+ICAM1-coated surfaces on TILs phenotype, we harvested the low- and high-expanding TILs at the end of the REP process (days 14) and subjected them to spectral flow cytometry using selected phenotypic markers. We found that the vast majority of the low-expanding TILs, independently of whether they were cultured on a CCL21+ICAM1 surface or not, were CD4^+^ T cells **(Figure 7a)**. Notably, the average frequency of CD4^+^ and CD8^+^ TIL populations was higher in cells incubated on coated surfaces compared to those cultured on uncoated surfaces (%CD4 cells on coated surfaces= 53.6 ± 0.8%, compared to 32.1 ± 2.1% on uncoated surfaces (*p*<0.001); %CD8 on coated surfaces =31.0 ± 0.9%, compared to 18.5 ± 1.8% on uncoated surfaces (*p*<0.01), **Figure 7a**). The remaining analyzed CD3^+^ TIL not within these single positive gates were predominantly double negative cells. In contrast, the high-expanding TILs, independently of whether they were treated with the SIN or not, were predominantly CD8^+^ T cells **(Figure 7e)**. In addiation, the prominance of CD4^+^ and CD8^+^ TIL populations within SIN-treated TILs was similar to that of cells cultured on uncoated surfaces (%CD4 cells on coated surfaces= 20.8 ± 4.2%, compared to 26.6 ± 5.9% on uncoated surfaces; %CD8 on coated surfaces = 82.2± 8.9%, compared to 70.9± 9.9% on uncoated surfaces, **Figure 7e**).

**Fig. 7:**
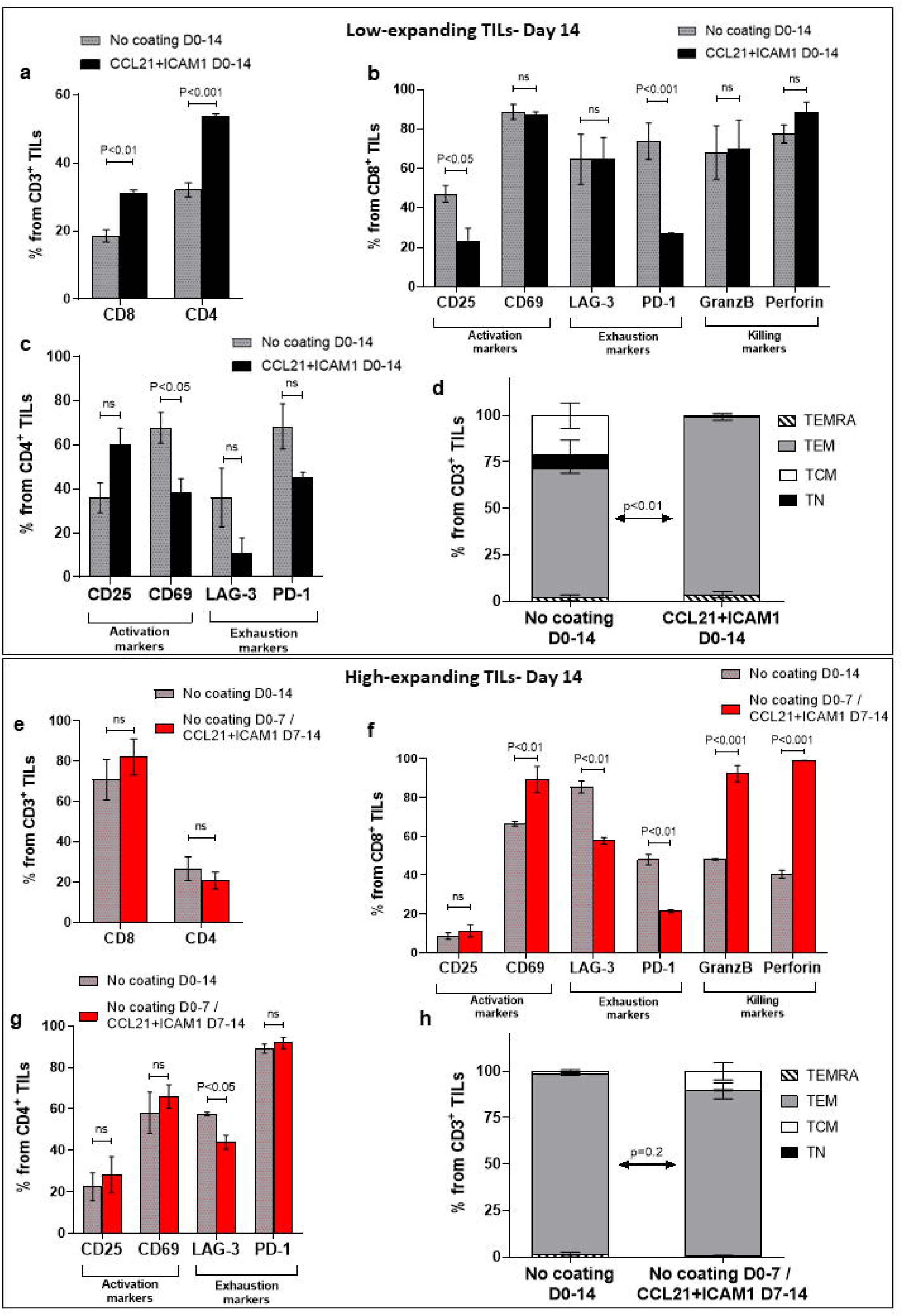
Molecular phenotyping of SIN-stimulated TILs at the end of the REP process (day 14). Low- and high-expanding TILs (upper panel **(a-d)** and lower panel **(e-h),** respectively) were harvested on day 14, at the end of the REP process and subjected to spectral flow cytometry using selected phenotypic markers. The low-expanding TILs underwent simultaneous treatment with feeder cells and SIN for 14 days, whereas the high-expanding TILs were treated for 7 days with feeder cells only, followed by SIN treatment, without feeder cells for an additional period of 7 days. Data are shown as mean ± SEM of three independent experiments. (**a** and **e)** Bar graphs illustrating the percentage of CD8^+^ and CD4^+^ T cells among the CD3^+^ T cells as determined by flow cytometry analysis. (**b, c, f, g)** Distribution of surface markers among CD8^+^ **(b, f)** and CD4^+^ T cells **(c, g),** that are related to T-cell activation (CD25, CD69), exhaustion (LAG-3, PD-1), and cytotoxic potency (GranzB, perforin) in low- **(b, c)** and high-expanding TILs **(f, g)**, in the presence or absence of SIN stimulation. (**d, h)** Distribution of differentiation subsets markers in low- **(d)** and high **(h)**-expanding TILs on day 14. The percentages (mean ± SEM) of naïve (TN, CD45RO^+^CCR7^+^), central memory (TCM, CD45RO^−^CCR7^+^), effector memory (TEM, CD45RO^−^CCR7^−^) and effector (TEMRA, CD45RO^+^CCR7^−^) T cells in CD3 population are represented. The data shown here are representatives of three independent experiments. Calculated p-values (using standard t-test) between the proportion of effector memory T cell subset in SIN-treated cells compared to untreated cells are as indicated in the figure.

Next, we evaluated the prominence of cells expressing specific phenotypic markers within the low-and high-expanding cell populations. As shown in **Figure 7b**, compared to untreated cells, SIN-treated CD8^+^ TILs displayed decreased expression of CD25 and PD-1, whereas no significant difference was evident in the expression of CD69, LAG-3, granzyme B and perforin. In contrast, SIN-treated CD4^+^ TILs displayed decreased expression of CD69, whereas no significant difference was evident in the expression of CD25, LAG-3 and PD-1 **(Figure 7c)**. Analysis of the T_Naïve_ cell and memory T-cell subsets revealed that, compared with the control, substrate-immobilized CCL21 + ICAM1 substantially increased the prominence of effector memory T cells (TEM) in low-expanding TILs (*p*<0.01) **(Figure 7d)**. Taken together, these results strongly suggest that the contribution of CCL21+ICAM1 SIN to the elevated cell yield of low-expanding TILs was due to decreased in expression of exhaustion related markers.

Analysis of the high-expanding TILs revealed that SIN-treated CD4^+^ TILs did not show significant differences in the expression levels of CD25 and CD69, compare to untreated cells (**Figure 7g).** However, compared with T cells cultured on uncoated surfaces, treatment of CD8^+^ TILs with SIN elevated the expression of CD69, whereas no significant difference was evident in the expression of CD25 **(Figure 7f)**. Notably, similar to that in low-expanding TILs, SIN treatment suppressed the expression of the inhibitory molecule PD-1 on CD8^+^ T cells (**Figure 7f),** but had no effect on the expression levels of PD-1 on CD4^+^ T cells **(Figure 7g)**. Additionally, SIN treatment reduced the expression of LAG-3 on both CD8^+^ and CD4^+^ T cells (**Figures 7f** and **g)**. Unlike the effects of SIN on low-expanding TILs, similar treatment of high-expanding TILs resulted in elevation of granzyme B and perforin expression on CD8^+^ T cells **(Figure 7f)**. Our data also revealed that the majority of high-expanding TILs, independently of whether they were cultured on a CCL21+ICAM1 surface or not, were effector memory T cells (*p*=0.2) **(Figure 7h)**. These results together suggest that stimulation of high-expanding TILs with CCL21+ICAM1 SIN greatly enhances the expression of killing markers and suppresses the expression of exhaustion related markers in these cells.

## Discussion

The present study addresses a key double challenge in current adoptive cancer immunotherapy, namely the need for both large numbers of autologous cancer-reactive cells and, at the same time, retention of high cytotoxic potency the these cells, ^14,47^ overcoming the natural tendency of T-cells to undergo exhaustion or anergy upon their *ex vivo* expansion. ^17,19^ To address this challenge, we have developed a “synthetic immune niche” (SIN) that apparently optimizes the balance between T-cell expansion and functionality. In our earlier studies, we have demonstrated that co-stimulation of activated murine CD4^+^ ^23^ and CD8^+ 22^ T cells with substrate-immobilized CCL21+ ICAM1 SIN, augments the expansion of both T-cell populations and further enhances the intrinsic cytotoxic activity of CD8^+^ cells both in culture and *in vivo.* ^22^ We further characterized the cellular and molecular processes associated with the capacity of the CCL21+ICAM1-based SIN to induce an “optimal interplay” between the proliferation and cytotoxicity of CD8^+^ T cells. ^35^ More recently, we showed that the CCL21+ICAM1 SIN can increase the expansion rate of melanoma patient-derived TILs, yet these experiments also revealed considerable variability in the REP expansion yields of the tested TILs, both in the presence and absence of the SIN. ^36^

In the present study, we addressed the basis for the patient-to-patient heterogeneity in TIL expansion by conducting an in-depth comparative phenotyping of diverse pairs of TILs and feeder cells. We show here that within the tested samples, high-expanding and low-expanding TILs with differential responsiveness to the SIN treatment could be clearly distinguished. We further demonstrated that the high- and low-expanding phenotypes can be predicted right after the pre-REP stage, based on morphological and molecular phenotyping, enabling the use of differential SIN stimulation protocols that enhance the proliferation of both the high- and the low-expanding TILs.

The results presented in **Figure 1a** clearly demonstrate two non-overlapping populations of pre-REP TILs: low-expanding cells (with typical fold expansion values of 68.3±5.8) and high-expanding TILs (typical fold expansion values of 244.8±10.2). Notably, these observations agree with a vast literature, pointing to major patient-to-patient differences in TIL expansion rates. ^44–46^ Importantly, at the end of the preREP stage, the low- and high-expanding TILs displayed, distinct morphological properties determined by flow cytometry and quantitative light microscopy imaging, as well as a differential profile of surface markers. Specifically, the high-expanding TILs displaying relatively high FSC and SSC values (by flow cytometry, **Figure 2b** and **b’**) or projected cell area above 120 µm^2^ (determined by oblique-optical microscopy, **Figure 1b** and **b’**). In our earlier study conducted with murine T cells, we have shown that T cells with high projected area display elevated proliferation and killing potency. ^35^

Further molecular profiling of pre-REP TILs, based on spectral flow cytometry analyses, revealed differences that may predict clinical responses. Notably, a low prominence of CD8^+^ T cells and high frequency of CD4^+^ T cells (i.e., a lower CD8/CD4 ratio) within the pre-REP TILs correlates with the high expansion levels of these cells **(Figure 2c** and **d)**. Importantly, our data also indicate that the prominence of CD4^−^ CD8^−^ subset is greater for low-expanding than for high-expanding TILs **(Figure 2c)**. These “double negative” T cells, which are a unique T cell subgroup, defined by the expression of CD3 with the absence of CD4, CD8 and CD56 cell markers, ^53,54^ have been shown to infiltrate solid tumors such as NSCLC, ^55^ liver cancer, ^56^ glioma, ^57^ and pancreatic tumors. ^58^ However, the effect of these double negative TILs in solid tumors remains largely unexplored, possibly due to their poor expansion capacity, as observed in our current study **(Figure 2c)**. Interestingly, previous evidence suggests that stimulation of CD4^+^ T cells *in vitro* for long periods (3 weeks) results in the generation of double negative T cells, ^59^ indicating that high prominence of the CD4^−^ CD8^−^ subset in low-expanding pre-REP TILs may be related to down-regulation of the CD4 molecule on these cells **(Figure 2c)**.

Our data further reveal that the low-expanding pre-REP TILs demonstrate a significant increase in LAG-3 expression, on both CD8^+^ **(Supplementary Fig. 3e)** and CD4^+^ **(Supplementary Fig. 3g)** T cells, compared the high-expanding pre-REP TILs. LAG-3 is associated with reduced proliferative capacity, ^42,43^ suggesting that this molecule attenuate the proliferation of the low-expanding TILs. Finally, we found that the majority of both low- and high-expanding TILs are classified as effector memory T cells (TEM: CD45RO^+^CCR7^−^, **Figure 2e**). Collectively, these results demonstrate that besides their distinct morphology, the high- and low-expanding pre-REP TILs can be readily distinguished by the relative prominence of CD4^+^, CD8^+^ and CD4^−^ CD8^−^ T cells, enabling the clear prediction of their REP expansion capacities.

Interestingly, when the same TILs underwent REP process with different feeder cell partners, consisting of a mixture of PBMC obtained from 2 healthy donors, different fold expansion values were obtained (**Figure 1a**). This additional complexity can be addressed by using feeder cells mixture of PBMC from three or more healthy donors, rather than two, which might compensate for occasional poor source of feeder cells and reduce the effect of variations between feeder cells from different donors on the expansion capacity of pre-REP TILs. ^60^

To address the differential effects of SIN treatment on the high- and low-expanding TILs, distinct REP processes were designed for the two TIL populations, in the presence or absence of the SIN. We found that the SIN treatment enhanced the expansion rate of the low-expanding TIL cultures (4.8-fold increase over cells cultured on the uncoated surface, **Figure 3 a, b** and **g).** On the other hand, SIN treatment of the high-expanding TILs revealed an unexpected loss of stimulatory effect of the SIN, or even suppression of TIL expansion **(Figure 3a-d)**, due to an apparent strong conflict between the concomitant feeder cell stimulation and SIN stimulation, manifested by a smaller yield of cells (**Figure 3i)**, compared to untreated cells (**Figure 3h**). Notably, elimination of feeder cells from the REP culture suppressed the TILs’ expansion, irrespective of the presence of SIN treatment. To explore the mechanism underlying the intriguing interplay between the feeder cells and the SIN, high-expanding pre-REP TILs were cultured with conditioned medium (CM) of feeder cells, in the presence or absence of the SIN. This experiment indicated that the CM partially supported TIL expansion in the absence of SIN, yet, it still suppressed the proliferation of TILs growing on the SIN **(Figure 4b** and **b’)**, indicating that the conflict is not induced by physical contact between the feeder cells and the T cells, but it is rather mediated by suppressing factors secreted to the medium by the feeder cells. Furthermore, replacement of the CCL21+ICAM1 SIN with surfaces coated with only one of the SIN components of the SIN **(Figure 4e)**, indicated that stimulation with CCL21 alone induced proliferation of TILs **(Figure 4c)**, while treatment with immobilized ICAM1 suppressed their proliferation **(Figure 4d)**.

Our attempts to identify, in the CM, specific components that might stimulate TILs (in the absence of SIN) and block TILs (in its presence) indicated that SIN treatment for 5 days strongly modulates the cytokine profile in feeder cells (alone) and feeder-TIL co-cultures **(Figure 5a** and **b**, and **Supplementary Figure 4a and b)**. Specifically, SIN increased the levels of CCL3/4, CXCL1, IL-1β, and IL-6 while suppressing IL-8 and CXCL10, suggesting a potential role in enhancing T-cell activation and persistence. These findings are consistent with previous studies showing that IL-6 and IL-1β support T-cell proliferation, ^49,50^ while IL-8 and CXCL10 contribute to T-cell dysfunction. ^47,48^ Notably, the observed upregulation of ICAM1, along with CCL3/4 and CXCL1, may further promote T-cell recruitment and activation, ^51,52^ potentially enhancing anti-tumor immune responses. These results highlight SIN’s potential to reshape the tumor immune microenvironment.

Taken together, these observations led to the development of the differential REP process, designed for the low- and high-expanding TILs, based on 14 days REP of the low expanding cells with feeders and SIN, and a temporal separation of feeder cells stimulation and the CCL21+CAM1 SIN treatment. Specifically, we found that co-culturing of the high-expanding TILs with the feeder cells for one-week, followed by one week with the SIN only, had a remarkable synergistic effect on TIL expansion (2.3-fold increase over cells cultured on the uncoated surface, **Figure 6e** and **f**).

To investigate the impact of CCL21+ICAM1 based SIN on TILs cytotoxic phenotype, the low-and high-expanding TILs were analyzed at the end of the REP, on day 14, for expression of selected phenotypic markers that were previously shown by us to be associated with high cytotoxic efficacy. ^35^ Specifically, we showed in that study, that murine T cells activated by antigen-loaded dendritic cells or by bead-conjugated into, CD3/CD28, and treated with SIN, reached optimal balance between proliferation and cytotoxic potency upon upregulation of cytotoxic gene signatures (following antigen-specific activation), and downregulation of exhaustion and proapoptotic genes optimizes the proliferation-cytotoxicity balance in the bead-activated cells. ^35^

A similar profiling of the low-expanding TILs, independently of whether they were treated with the SIN or not, indicated that these cells were predominantly CD4^+^ T cells **(Figure 7a)**, similar to what was observed in the pre-REP high-expanding TILs (**Figure 2c).** Of note, the prominence of CD4^+^ and CD8^+^ TIL subsets was higher in SIN-treated cells compared to those cultured on uncoated surfaces (**Figure 7a**). In contrast, the high-expanding TILs, irrespective of the SIN treatment, were predominantly CD8^+^ T cells **(Figure 7e)**. Addiationaly, SIN treatment had no effect on the average frequency of CD4^+^ and CD8^+^ cells in these TILs (**Figure 7e**). Adoptive cancer immunotherapy efficacy depends both quantity and quality of the expanded lymphocytes. Predominantly CD8^+^-rich products generally correlate with better antitumor activity *in vitro* ^61^ and *in vivo*; ^62,63^ though, transferred CD4^+^ TILs also exhibit antitumor effects and can lead to clinical responses. ^64,65^

Phenotypic profiling of the low-expanding cells on day 14, indicated suppressed expression of PD1, in the SIN-treated CD8^+^ cells, and high levels of expression of granzyme B and perforin. The high-expanding TILs, following 7 days co-culturing with feeder cells, and additional 7 days with the CCL21+ICAM1 SIN displayed a strong suppression of both PD-1 and LAG-3, and pronounced elevation of granzyme B and perforin. Collectively, these data strongly suggest that the CCL21+ICAM1 SIN enhances both the proliferation and cytotoxic potency of the low-expanding, as well as the high-expanding TILs, by suppressing the expression of exhaustion-related markers, and upregulating the prominence of granzyme and perforin in the high-expanding cells. Based on these results we propose that the novel REP process described here might greatly enhance the efficacy of TIL-based cancer immunotherapy.

## Conclusion

In this study, we addressed the effect of SIN treatment on the expansion of pre-REP tumor infiltrating lymphocytes. Specifically, we showed here that the pre-REP T-cells are heterogeneous in their proliferation capacity, with two distinct sub-populations, displaying either high- or low-expansion capacity. These two sub-populations display distinct responsiveness to the SIN stimulation; the low-expansion cells undergo major augmentation of their proliferation when cultured on the CCL21+ICAM1 SIN together with feeder cells for 2 weeks, while the high-expansion cells need a 7-day stimulation with feeder cells only, followed by 7-day incubation with the SIN, in the absence of feeders. The structural and molecular profiling of the pre-REP cells indicated that the high- and low-expanding TILs can be readily distinguished, enabling the application of the optimal SIN treatment for the cells. These results strongly suggest that incorporation of SIN stimulation into the TIL expansion process might significantly improve the therapeutic performance of these cells.

## Acknowledgments

We would like to thank Yoseph Addadi and Inna Goliand from the de Picciotto Cancer Cell Observatory (WIS), and Ekaterina Kopitman and the Flow Cytometry Unit (WIS) for their expert technical assistance.

## Authors’ Contributions

Conceptual and experimental design: BG, MB, SY, RZ, MM

Performance of the experiments and resources: SY, RZ, SA, TU, KB, AN, AG

Data analysis and quantification: SY, RZ

Manuscript writing and approval: SY, RZ, MB, BG

## References

1. Tan, S., Li, D. & Zhu, X. Cancer immunotherapy: Pros, cons and beyond. Biomed. Pharmacother. 124, (2020).

2. Rohaan, M. W., Wilgenhof, S. & Haanen, J. B. A. G. Adoptive cellular therapies: the current landscape. Virchows Arch. 474, 449–461 (2019).

3. Bear, A. S., Fraietta, J. A., Narayan, V. K., O’Hara, M. & Haas, N. B. Adoptive Cellular Therapy for Solid Tumors. Am. Soc. Clin. Oncol. Educ. B. 57–65 (2021) doi:10.1200/edbk_321115.

4. Parsonidis, P. & Papasotiriou, I. Adoptive Cellular Transfer Immunotherapies for Cancer. Cancer Treat. Res. Commun. 32, 100575 (2022).

5. Creelan, B. C. et al. Tumor-infiltrating lymphocyte treatment for anti-PD-1-resistant metastatic lung cancer: a phase 1 trial. Nat. Med. 27, 1410–1418 (2021).

6. Stevanović, S. et al. Complete regression of metastatic cervical cancer after treatment with human papillomavirus-targeted tumor-infiltrating T cells. J. Clin. Oncol. 33, 1543–1550 (2015).

7. Tran, E. et al. T-Cell Transfer Therapy Targeting Mutant KRAS in Cancer. N. Engl. J. Med. 375, 2255–2262 (2016).

8. Zacharakis, N. et al. Immune recognition of somatic mutations leading to complete durable regression in metastatic breast cancer. Nat. Med. 24, 724–730 (2018).

9. Rosenberg, S. A. et al. Durable complete responses in heavily pretreated patients with metastatic melanoma using T-cell transfer immunotherapy. Clin. Cancer Res. 17, 4550–4557 (2011).

10. Goff, S. L. et al. Randomized, prospective evaluation comparing intensity of lymphodepletion before adoptive transfer of tumor-infiltrating lymphocytes for patients with metastatic melanoma. J. Clin. Oncol. 34, 2389–2397 (2016).

11. Besser, M. J. et al. Clinical responses in a phase II study using adoptive transfer of short-term cultured tumor infiltration lymphocytes in metastatic melanoma patients. Clin. Cancer Res. 16, 2646–2655 (2010).

12. Kwong, M. L. M. & Yang, J. C. Lifileucel : FDA-approved T-cell therapy for melanoma. 2, 648–650 (2024).

13. Hulen, T. M., Chamberlain, C. A., Svane, I. M. & Met, Ö. ACT Up TIL Now: The Evolution of Tumor-Infiltrating Lymphocytes in Adoptive Cell Therapy for the Treatment of Solid Tumors. Immuno 1, 194–211 (2021).

14. Granhøj, J. S. et al. Tumor-infiltrating lymphocytes for adoptive cell therapy: recent advances, challenges, and future directions. Expert Opin. Biol. Ther. 22, 627–641 (2022).

15. Morotti, M. et al. Promises and challenges of adoptive T-cell therapies for solid tumours. Br. J. Cancer 124, 1759–1776 (2021).

16. Peterson, C., Denlinger, N. & Yang, Y. Recent Advances and Challenges in Cancer Immunotherapy. Cancers (Basel). 14, (2022).

17. Taefehshokr, S. et al. Cancer immunotherapy: Challenges and limitations. Pathol. Res. Pract. 229, 153723 (2022).

18. Yang, L., Ning, Q. & Tang, S. S. Recent Advances and Next Breakthrough in Immunotherapy for Cancer Treatment. J. Immunol. Res. 2022, (2022).

19. Waldman, A. D., Fritz, J. M. & Lenardo, M. J. A guide to cancer immunotherapy: from T cell basic science to clinical practice. Nat. Rev. Immunol. 20, 651–668 (2020).

20. Van Den Berg, J. H., et al. Tumor infiltrating lymphocytes (TIL) therapy in metastatic melanoma: Boosting of neoantigen-specific T cell reactivity and long-term follow-up. J. Immunother. Cancer 8, 1–11 (2020).

21. Brown, L. V., Gaffney, E. A., Ager, A., Wagg, J. & Coles, M. C. Quantifying the limits of CAR T-cell delivery in mice and men. J. R. Soc. Interface 18, (2021).

22. Adutler-Lieber, S., Friedman, N. & Geiger, B. Expansion and antitumor cytotoxicity of T-Cells are augmented by substrate-bound CCL21 and intercellular adhesion molecule 1. Front. Immunol. 9, (2018).

23. Adutler-Lieber, S. et al. Substrate-bound CCL21 and ICAM1 combined with soluble IL-6 collectively augment the expansion of antigen-specific murine CD41 T cells. Blood Adv. 1, 1016–1030 (2017).

24. Luo, H. et al. Coexpression of IL7 and CCL21 increases efficacy of CAR-T cells in solid tumors without requiring preconditioned lymphodepletion. Clin. Cancer Res. 26, 5494–5505 (2020).

25. Willimann, K. et al. The chemokine SLC is expressed in T cell areas of lymph nodes and mucosal lymphoid tissues and attracts activated T cells via CCR7. Eur. J. Immunol. 28, 2025–2034 (1998).

26. Gunn, M. D. et al. A chemokine expressed in lymphoid high endothelial venules promotes the adhesion and chemotaxis of naive T lymphocytes. Proc. Natl. Acad. Sci. U. S. A. 95, 258–263 (1998).

27. Dikovsky, D., Bianco-Peled, H. & Seliktar, D. The effect of structural alterations of PEG-fibrinogen hydrogel scaffolds on 3-D cellular morphology and cellular migration. Biomaterials 27, 1496–1506 (2006).

28. Friedman, R. S., Jacobelli, J. & Krummel, M. F. Surface-bound chemokines capture and prime T cells for synapse formation. Nat. Immunol. 7, 1101–1108 (2006).

29. Flanagan, K., Moroziewicz, D., Kwak, H., Hörig, H. & Kaufman, H. L. The lymphoid chemokine CCL21 costimulates naïve T cell expansion and Th1 polarization of non-regulatory CD4+ T cells. Cellular Immunology vol. 231 75–84 at 10.1016/j.cellimm.2004.12.006 (2004).

30. Molon, B. et al. T cell costimulation by chemokine receptors. Nat. Immunol. 6, 465–471 (2005).

31. Gollmer, K. et al. CCL21 mediates CD4+ T-cell costimulation via a DOCK2/Rac-dependent pathway. Blood 114, 580–588 (2009).

32. Ingulli, E., Mondino, A., Khoruts, A. & Jenkins, M. K. In vivo detection of dendritic cell antigen presentation to CD4+ T cells. J. Exp. Med. 185, 2133–2141 (1997).

33. Woolf, E. et al. Lymph node chemokines promote sustained T lymphocyte motility without triggering stable integrin adhesiveness in the absence of shear forces. Nat. Immunol. 8, 1076–1085 (2007).

34. Stein, J. V. et al. The CC chemokine thymus-derived chemotactic agent 4 (TCA-4, secondary lymphoid tissue chemokine, 6Ckine, exodus-2) triggers lymphocyte function-associated antigen 1-mediated arrest of rolling T lymphocytes in peripheral lymph node high endothelial venule. J. Exp. Med. 191, 61–75 (2000).

35. Yado, S. et al. Molecular mechanisms underlying the modulation of T-cell proliferation and cytotoxicity by immobilized CCL21 and ICAM1. J. Immunother. cancer 12, 1–18 (2024).

36. Yunger, S., Geiger, B. & Friedman, N. Modulating the proliferative and cytotoxic properties of patient-derived TIL by a synthetic immune niche of immobilized CCL21 and ICAM1. 1–10 (2023) doi:10.3389/fonc.2023.1116328.

37. Besser, M. J. et al. Adoptive transfer of tumor-infiltrating lymphocytes in patients with metastatic melanoma: Intent-to-treat analysis and efficacy after failure to prior immunotherapies. Clin. Cancer Res. 19, 4792–4800 (2013).

38. Stringer, C., Wang, T., Michaelos, M. & Pachitariu, M. Cellpose: a generalist algorithm for cellular segmentation. Nat. Methods 18, 100–106 (2021).

39. Fenton, G. A. & Mitchell, D. A. Cellular Cancer Immunotherapy Development and Manufacturing in the Clinic. Clin. Cancer Res. 29, 843–857 (2023).

40. Li, B. Why do tumor-infiltrating lymphocytes have variable efficacy in the treatment of solid tumors? Front. Immunol. 13, 1–13 (2022).

41. Blank, C. U. et al. Defining ‘T cell exhaustion’. Nat. Rev. Immunol. 19, 665–674 (2019).

42. Ma, J. et al. Blockade of PD-1 and LAG-3 expression on CD8+ T cells promotes the tumoricidal effects of CD8+ T cells. Front. Immunol. 14, 1–13 (2023).

43. Lichtenegger, F. S. et al. Targeting LAG-3 and PD-1 to enhance T cell activation by antigen-presenting cells. Front. Immunol. 9, 1–12 (2018).

44. Poschke, I. C. et al. The Outcome of Ex Vivo TIL Expansion Is Highly Influenced by Spatial Heterogeneity of the Tumor T-Cell Repertoire and Differences in Intrinsic in Vitro Growth Capacity between T-Cell Clones. Clin. Cancer Res. 26, 4289–4301 (2020).

45. Knochelmann, H. M. et al. Modeling ex vivo tumor-infiltrating lymphocyte expansion from established solid malignancies. Oncoimmunology 10, 1–10 (2021).

46. Kazemi, M. H. et al. Tumor-infiltrating lymphocytes for treatment of solid tumors: It takes two to tango? Front. Immunol. 13, 1–23 (2022).

47. David, J. M., Dominguez, C., Hamilton, D. H. & Palena, C. The IL-8/IL-8R axis: A double agent in tumor immune resistance. Vaccines 4, (2016).

48. Liu, M., Guo, S. & Stiles, J. K. The emerging role of CXCL10 in cancer. Oncol. Lett. 2, 583–589 (2011).

49. Hirano, T. IL-6 in inflammation, autoimmunity and cancer. Int. Immunol. 33, 127–148 (2021).

50. Shiomi, A., Usui, T. & Mimori, T. GM-CSF as a therapeutic target in autoimmune diseases. Inflamm. Regen. 36, 1–9 (2016).

51. Oo, Y. H., Shetty, S. & Adams, D. H. The role of chemokines in the recruitment of lymphocytes to the liver. Dig. Dis. 28, 31–44 (2010).

52. Griffith, J. W., Sokol, C. L. & Luster, A. D. Chemokines and chemokine receptors: Positioning cells for host defense and immunity. Annu. Rev. Immunol. 32, 659–702 (2014).

53. Wu, Z. et al. CD3+CD4-CD8-(Double-Negative) T Cells in Inflammation, Immune Disorders and Cancer. Front. Immunol. 13, 1–14 (2022).

54. Velikkakam, T., Gollob, K. J. & Dutra, W. O. Double-negative T cells: Setting the stage for disease control or progression. Immunology 165, 371–385 (2022).

55. Fang, L. et al. Targeting late-stage non-small cell lung cancer with a combination of DNT cellular therapy and PD-1 checkpoint blockade. J. Exp. Clin. Cancer Res. 38, 1–14 (2019).

56. Di Blasi, D. et al. Unique T-Cell Populations Define Immune-Inflamed Hepatocellular Carcinoma. Cmgh 9, 195–218 (2020).

57. Liu, Z. et al. Tumor-infiltrating lymphocytes (TILs) from patients with glioma. Oncoimmunology 6, (2017).

58. Hall, M. L. et al. Expansion of tumor-infiltrating lymphocytes (TIL) from human pancreatic tumors. J. Immunother. Cancer 4, 1–12 (2016).

59. Grishkan, I. V., Ntranos, A., Calabresi, P. A. & Gocke, A. R. Helper T cells down-regulate CD4 expression upon chronic stimulation giving rise to double-negative T cells. Cell. Immunol. 284, 68–74 (2013).

60. Hopewell, E. L., Cox, C., Pilon-Thomas, S. & Kelley, L. L. Tumor-infiltrating lymphocytes: Streamlining a complex manufacturing process. Cytotherapy 21, 307–314 (2019).

61. Schwartzentruber, D. J. et al. In vitro predictors of therapeutic response in melanoma patients receiving tumor-infiltrating lymphocytes and interleukin-2. J. Clin. Oncol. 12, 1475–1483 (1994).

62. Radvanyi, L. G. et al. Specific lymphocyte subsets predict response to adoptive cell therapy using expanded autologous tumor-infiltrating lymphocytes in metastatic melanoma patients. Clin. Cancer Res. 18, 6758–6770 (2012).

63. Itzhaki, O. et al. Establishment and large-scale expansion of minimally cultured young tumor infiltrating lymphocytes for adoptive transfer therapy. J. Immunother. 34, 212–220 (2011).

64. Qui, H. Z. et al. CD134 Plus CD137 Dual Costimulation Induces Eomesodermin in CD4 T Cells To Program Cytotoxic Th1 Differentiation. J. Immunol. 187, 3555–3564 (2011).

65. Antony, P. A. et al. CD8+ T Cell Immunity Against a Tumor/Self-Antigen Is Augmented by CD4+ T Helper Cells and Hindered by Naturally Occurring T Regulatory Cells. J. Immunol. 174, 2591–2601 (2005).

