## Supplementary information for "Differential effects of immobilized CCL21 and ICAM1 on TILs with distinct expansion properties"

**Supplementary Figures**

**Supplementary Fig. 1**

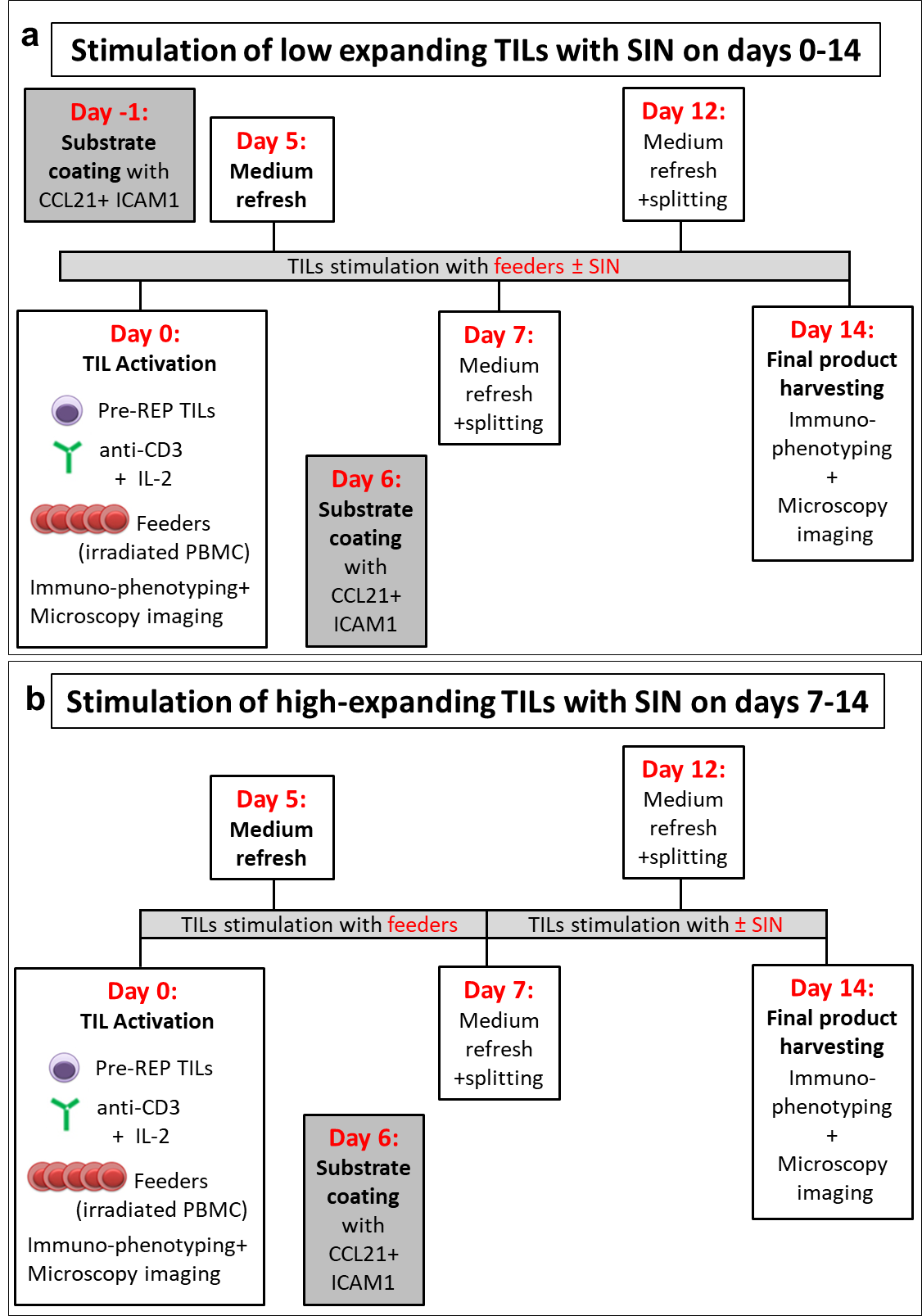

**Supplementary Fig. 1: A schematic representation of a 14-days REP process with CCL21 and ICAM1 surface coating.** To determine the differential effects of SIN treatment on the high- and low-expanding TILs, both populations of cells were stimulated with anti-CD3 mAb, IL-2 and irradiated feeder cells, in the presence or absence of SIN stimulation. The low-expanding TILs were cultured on CCL21 + ICAM1 substrates at the initiation of the REP (Day 0, **a**), while the high-expanding TILs were exposed to CCL21+ICAM1 on days 7-14 of the REP (**b**).

**Supplementary Fig. 2**

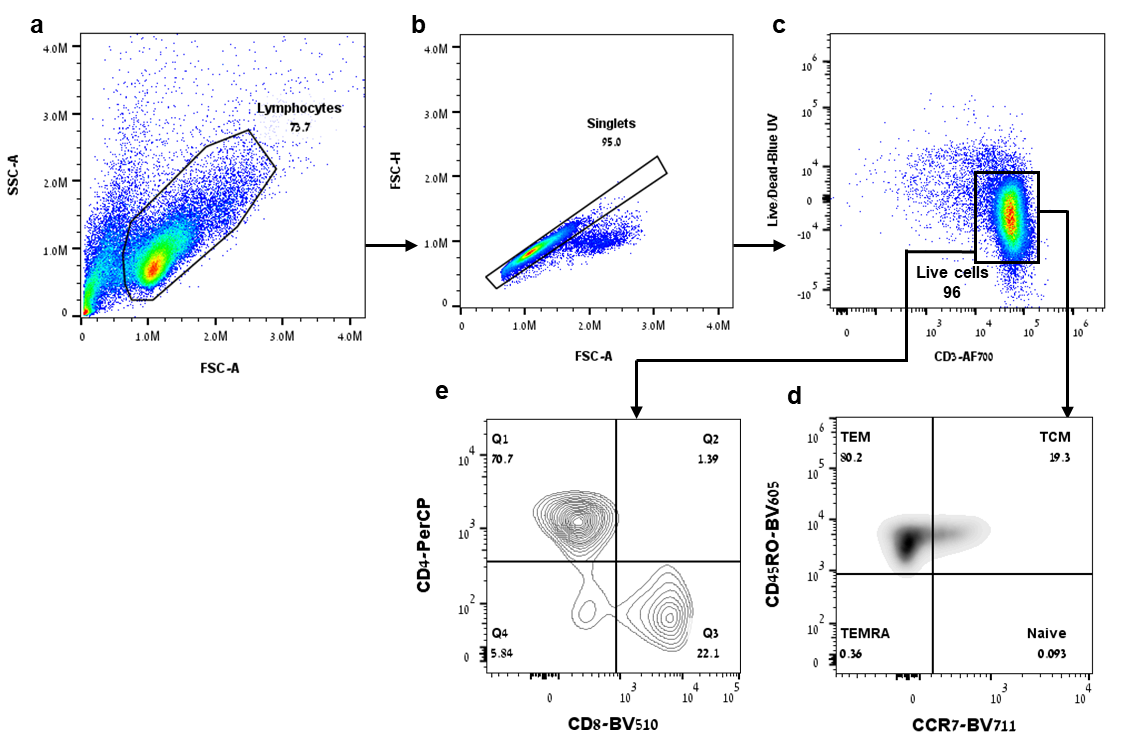

**Supplementary Fig. 2**: A representative gating strategy used for the phenotyping of T cells. (a) Total lymphocytes were first gated on a forward (FS)/side (SS) scatter plot, and then doublets were eliminated (b). Live CD3^+^ cells were selected (c) and were further analyzed for the expression of CD45RO in combination with CCR7 (d) and CD4 in combination with CD8 (e). The CD4^+^ and CD8^+^ subsets were subsequently assessed for the expression of activation and exhaustion markers, including CD25, CD69, PD-1, LAG-3, and CD8^+^ population was further analyzed for the expression of the cytotoxic effector molecules Granzyme B and Perforin.

**Supplementary Fig. 3**

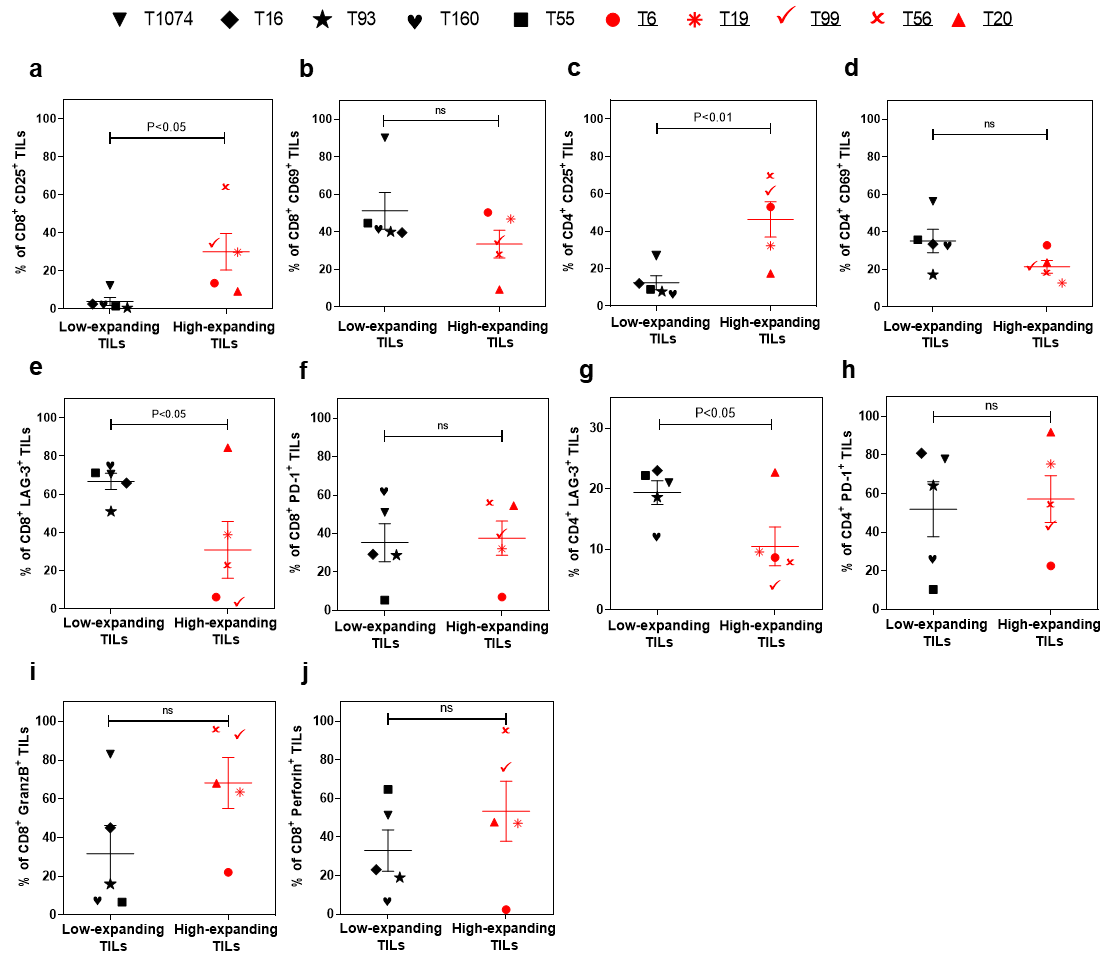

**Supplementary Fig. 3: Molecular phenotyping of pre-REP TILs enables a differential prediction of their expansion potency.** Flow cytometry-based phenotyping of pre-REP samples (n=10). Live CD8^+^ and CD4^+^ T cells were selected and analyzed for the expression of the following markers: CD25 (**a** and **c**), CD69 (**b** and **d**), LAG-3 (**e** and **g**), PD-1 (**f** and **h**), Granzyme B (**i)** and perforin (**j)**. Data are shown as mean ± SEM of three independent experiments. Calculated p-values (using standard t-test) are as indicated in the figure.

**Supplementary Fig. 4**

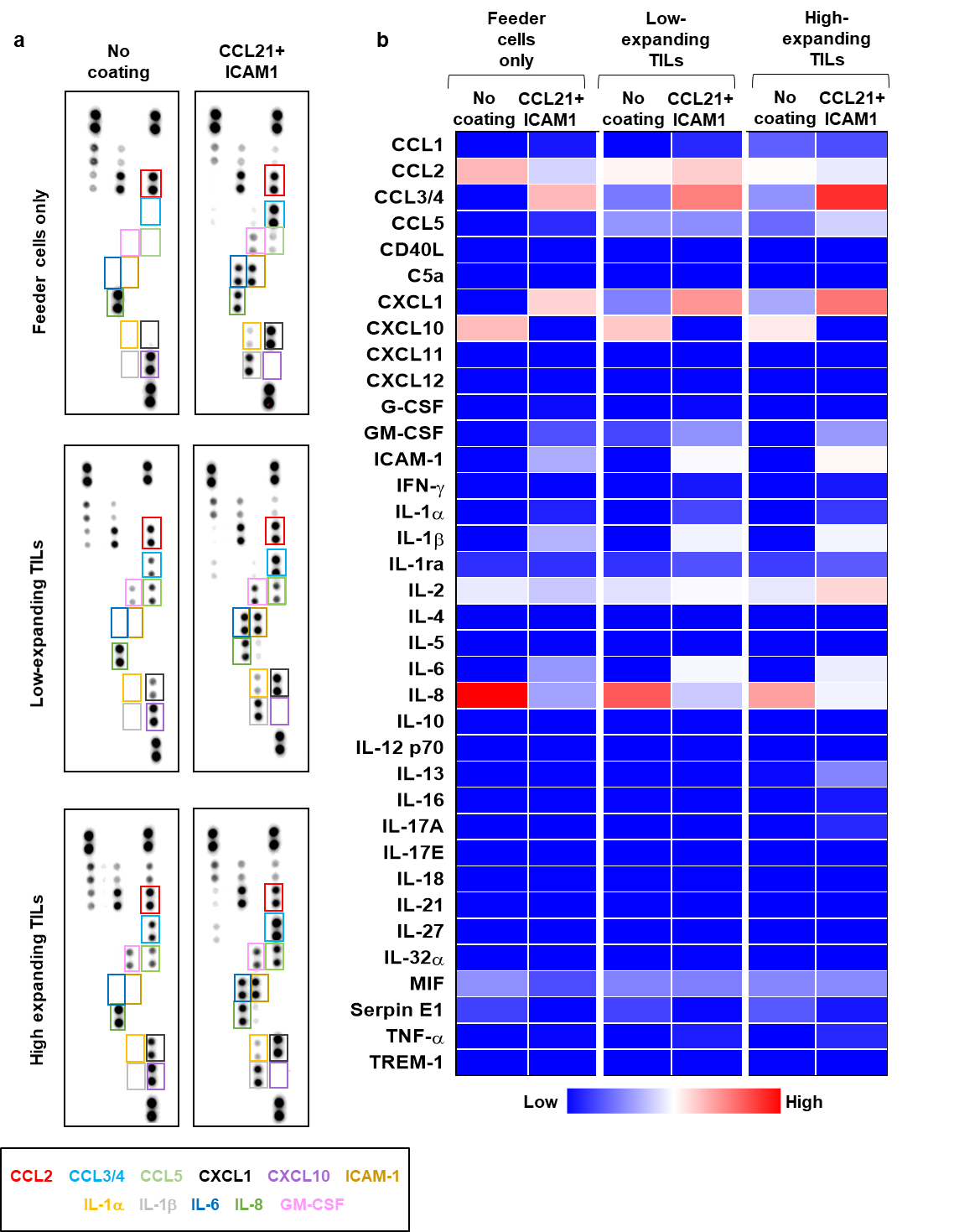

**Supplementary Fig. 4: The differential effects of CCL21+ICAM1 SIN treatment on cytokine secretion by low- and high-expanding TILs.**

Cytokine array analysis of culture supernatants collected at day 5 of the REP from feeder cells cultured alone (F_3_) or low- and high-expanding TIL cultures stimulated with feeder cells (T99/F_3_ and T55/F_3_). **(a)** Representative images for cytokine array panels. **(b)** Quantification of results from the cytokine array analysis. Heatmap represents average values from two dots in the array. The differentially secreted cytokines are highlighted in boxes and their labels are showed below with corresponding colors. Results of a repeated analysis with feeder T2 are shown in figure 5.

**Supplementary Methods**

Production and purification of human CCl21 (hCCL21):

The gene encoding hCCL21 (24-134) followed by a TEV cleavage site, and a C-terminal 6 x His tag was cloned into pET21a. The resulting construct was transformed into *E. coli* Bl21 (DE3*)* competent cells. Bacterial pre-cultures were grown overnight at 37°C with shaking and used to inoculate 7.5L LB. Cultures were grown at 37°C with shaking to an optical density OD_600_= 0.6. Protein expression was induced by the addition of isopropyl-β-d-thiogalactopyranoside (IPTG; 200 μM), and the culture was kept at 15°C with shaking overnight. Cells were harvested by centrifugation, and the pellet was resuspended in 100 ml lysis buffer containing 50mM Tris pH 8, 300mM NaCl,10mM Imidazole,1mM DTT supplemented with 0.2mg/ml lysozyme, 20ug/ml DNase, 1mM MgCl_2_, and protease inhibitor cocktail (Calbiochem set 3). After lysis by a cooled protein disrupter (Constant Systems), the soluble fraction was obtained by centrifugation and purified by immobilized metal ion affinity chromatography (IMAC) using a HiTrap *FF_*5ml cartridge (Cytiva) using an FPLC system (ÄKTA GE Healthcare Life Sciences) at 4^o^C. The column was washed with lysis buffer containing 50mM imidazole and hCCl21 was eluted in one step with elution buffer (50mM Tris pH 8, 300mM NaCl, 500mM imidazole). The fractions containing hCCl21 were analyzed using SDS-PAGE, pooled and dialyzed into PBS overnight. hCCl21 was applied to an Xbridge C4 (Waters) reverse phase column operated by an HPLC system (Waters) pre-equilibrated with binding buffer containing 5% acetonitrille, 0.1%TFA in DDW. The protein was eluted with a linear gradient to 95% acetonitrille, 0.1%TFA in DDW. Pure hCCl21 samples from consecutive runs were pooled and lyophilize to dryness. The dry protein was resuspended in DDW and divided into small aliquots before flash freezing in liquid nitrogen.

Expression and purification of human ICAM1 (hICAM1):

hICAM1 (aa residues 28-480) followed by an Fc domain, a TEV recognition site, and a C-terminal 6 x His tag was cloned into pHLSec vector for secretion from Expi293 human cells. Medium (300ml) containing the secreted protein was collected 72 hr. post transfection for purification.  After carefully removing the cells, the medium was applied to a Ni column (HisTrap_FF_5ml) equilibrated with PBS and the bound hICAM was washed with PBS containing 50mM imidazole and eluted with PBS containing 500mM imidazole. Fractions containing hICAM1 were pooled, concentrated and injected to a Superdex_200_16/60_PrepGrad size exclusion column equilibrated with PBS. The identity of the protein was confirmed by SDS PAGE developed with Coomassie blue. Aliquots were flash frozen with liquid nitrogen and kept at -80^o^C.
